# Apical polarity and actomyosin dynamics regulate Hippo signaling by controlling Kibra subcellular localization

**DOI:** 10.1101/2023.06.05.543736

**Authors:** Sherzod A. Tokamov, Nicki Nouri, Ashley Rich, Stephan Buiter, Michael Glotzer, Richard G. Fehon

**Affiliations:** Department of Molecular Genetics and Cell Biology; Committee on Development, Regeneration, and Stem Cell Biology The University of Chicago, Chicago, IL 60637, USA; Department of Cell Biology, Duke University School of Medicine, Durham, NC, USA

## Abstract

Cell polarity and actomyosin networks are key features that organize cells of epithelial tissues and are known to regulate tissue growth. However, the mechanisms by which polarity cues and actomyosin cytoskeleton influence intracellular signaling cascades that control growth remain poorly understood. The Hippo pathway is an evolutionarily conserved regulator of tissue growth and is known to integrate inputs from both polarity and actomyosin components. An upstream activator of the Hippo pathway, Kibra, localizes into distinct pools at the junctional and medial regions of the apical cortex in epithelial cells, and medial accumulation was shown to promote Kibra activity. Here, we demonstrate that cortical Kibra distribution is controlled by a tug of war between apical polarity and actomyosin dynamics. We show that while the apical polarity network, in part via aPKC, tethers Kibra at the junctional cortex to silence its activity, medial actomyosin flows promote Kibra-mediated Hippo complex formation at the medial cortex, thereby activating the Hippo pathway. This study provides a mechanistic understanding of the relationship between the Hippo pathway, polarity, and actomyosin cytoskeleton and offers novel insights into how fundamental features of epithelial tissue architecture can serve as inputs into signaling cascades that control tissue growth, patterning, and morphogenesis.

## Introduction

Growth and morphogenesis of epithelial tissues are critical determinants of organ size. Epithelial tissues such as the Drosophila wing imaginal disc are composed of cells polarized along the apico-basal axis and tightly interconnected by intercellular junctions along the lateral domain. These characteristics feature prominently in the organization and function of the Hippo pathway, an evolutionarily conserved regulator of tissue growth (Boggiano et al., 2011; Genevet and Tapon, 2011). For example, an upstream activator of the Hippo pathway, Expanded (Ex), localizes at the junctional cortex (a term we use to include the marginal zone and the adherens junctions; Tepass, 2012), where it assembles and activates a signaling cascade that includes Hippo (Hpo) and Warts (Wts) kinases. The kinase cascade represses a pro-growth transcriptional effector Yorkie (Yki, Boedigheimer et al., 1997; McCartney et al., 2000; Hamaratoglu et al., 2006). Unlike Ex, Kibra (Kib) and Merlin (Mer) accumulate at both the junctional and apicomedial cortex (referred to as the “medial” cortex in this study), where Kib assembles a Hippo signaling complex that inhibits Yki in parallel to Ex (Su et al., 2017; Tokamov et al., 2021; Wang et al., 2022). The functional significance of these distinct subcellular Kib pools and the mechanisms that control the distribution of Kib between the junctional and medial cortex remain elusive.

Apico-basal asymmetry in epithelial cells is regulated by a mutually antagonistic relationship between apical and basolateral polarity determinants (Tepass, 2012). Among the apical components, a transmembrane protein Crumbs (Crb) regulates polarity via associated cytoplasmic partners, including the Stardust (Sdt)/Patj and Par-6/atypical protein kinase C (aPKC) complexes. Aside from its role in polarity, however, Crb recruits Ex to the junctional cortex, thereby enabling Ex-mediated Hippo pathway activation (Ling et al., 2010; Robinson et al., 2010). In the absence of Crb, Ex is diffuse throughout the cytoplasm and inactive. Previous work suggested that Crb also tethers Kib at the junctional cortex by an unknown mechanism, which suppresses Kib function (Su et al., 2017). Additionally, ectopic aPKC activity was previously shown to inhibit Hippo signaling (Grzeschik et al., 2010). Conversely, loss of Hippo pathway activity, at least in part via Yki-mediated transcriptional activity, promotes expansion of the apical domain in imaginal epithelial cells (Genevet et al., 2009). Thus, apico-basal polarity and Hippo signaling are highly interconnected.

Another critical regulator of Hippo signaling is the actomyosin cytoskeleton. In many epithelia, the cortical actomyosin network is organized into junctional and medial pools at the apical domain (Miao and Blankenship, 2020). Junctional actomyosin cables can generate sustained forces to resist mechanical strain or drive morphogenetic events, such as junctional remodeling during convergent extension. On the other hand, the medial actomyosin network is more transient and pulsatile and is important in driving apical constriction of epithelial cells, which leads to tissue scale deformations. A large body of evidence suggests that increased F-actin polymerization and cortical tension generated by non-muscle myosin II (MyoII) contractility promote the activity of Yki and its mammalian homolog, YAP (Fernandez et al., 2011; Sansores-Garcia et al., 2011; Aragona et al., 2013; Rauskolb et al., 2014; Ibar et al., 2018). Additionally, in cultured cells cortical actin inhibits the ability of Mer to recruit and activate Wts (Yin et al., 2013). However, how actomyosin organization and dynamics influence upstream Hippo signaling components remains unknown. Specifically, the relationship between medial actomyosin pulses and Hippo signaling has not been explored.

Several observations suggest that Kib is uniquely poised to mediate interactions between polarity, actomyosin, and Hippo signaling. First, Kib has a key role in recruiting and activating the Hippo kinase cascade (Genevet et al., 2010; Yu et al., 2010; Baumgartner et al., 2010; Su et al., 2017). Second, previous studies have shown that Kib physically interacts with aPKC (Büther et al., 2004; Yoshihama et al., 2011; Jin et al., 2015). Lastly, Kib’s localization at the junctional and medial cortex closely resembles that of actomyosin (Fig. 1A, Su et al., 2017). Here, using a combination of osmotic, pharmacological, and genetic manipulations we show that Kib distribution at the cell cortex is regulated by the apical polarity and actomyosin networks. We demonstrate that aPKC tethers Kib at the junctional cortex to inhibit Kib-mediated Hippo pathway activation, whereas medial actomyosin dynamics promote Kib activity by accumulating and maintaining it at the medial cortex. Our results suggest a model whereby a tug of war between the apical polarity and actomyosin networks modulates Hippo signaling and tissue growth via Kib subcellular localization.

**Figure 1:**
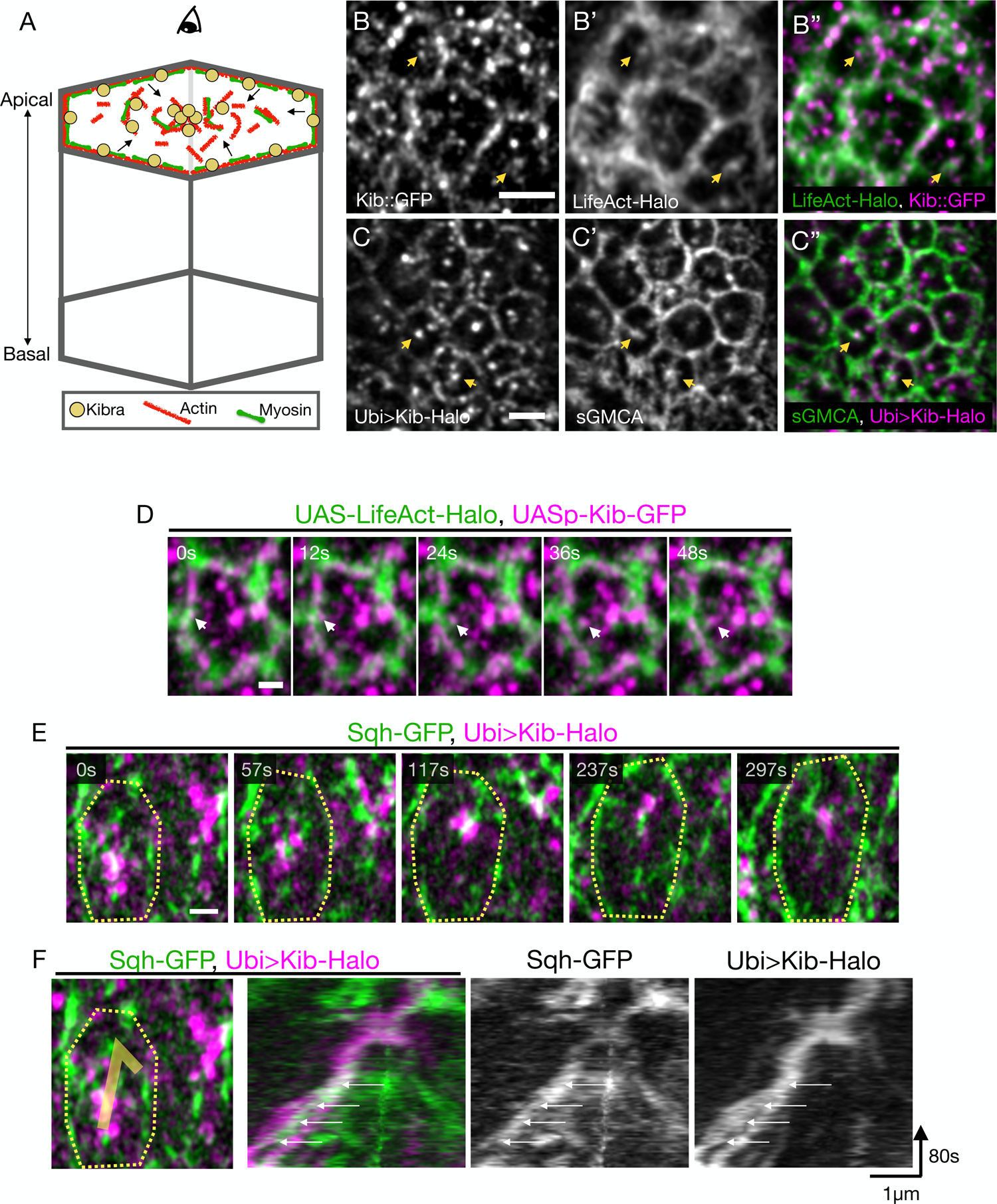
Medial Kib is associated with the actomyosin network. A) A cartoon of an epithelial cell polarized along the apical-basal axis. The actomyosin network and Kib are similarly partitioned at the apical cortex into junctional and medial pools. The arrows indicate flows of actomyosin components from the junctional to the medial region. Throughout the paper, images display tissue apical projections as viewed from the top. B-C’’) Apical projections of wing imaginal tissues expressing either endogenous Kib::GFP and LifeAct-Halo (B-B’’) or Ubi>Kib-Halo and sGMCA (C-C’’). Yellow arrowheads point at medial Kib punctae decorating apical F-actin structures. Scale bars = 2µm. D) Single frames from Movie 1 of a single cell showing co-migration of a Kib cluster (magenta) with F-actin (green) from the junctional to the medial region (white arrow). Scale bar = 1µm. E) Single frames from Movie 2 of a single cell showing co-migration of medial Kib (magenta) with Sqh (green). The contour line roughly outlines the cell. Scale bar = 1µm. F) A kymograph tracking co-migration of medial Kib and Sqh shown in E. The white arrows point to a myosin pulse increasing in intensity over time that is associated with the merger of two initially separate Kib punctae.

## Results

### Medial Kib is associated with the actomyosin network

The actomyosin cytoskeleton is known to regulate growth via Yki, but the link between actomyosin dynamics and the upstream Hippo pathway components remains unknown (throughout the study, we use the term “dynamics” to describe general movement of actomyosin components at the cell cortex). The similarity in Kib and actomyosin subcellular distribution prompted us to investigate their localization in more detail. Using Airyscan confocal microscopy for better spatial resolution, we first examined the localization of either endogenous Kib tagged with the green fluorescent protein (Kib::GFP) or Halo-tagged Kib expressed under the ubiquitin promoter (Ubi>Kib-Halo) with respect to F-actin. To visualize F-actin, we used either UAS-driven LifeAct-Halo or a GFP-tagged Moesin actin-binding domain expressed under the *spaghetti squash* promoter (sGMCA). In the wing imaginal epithelium, both Kib::GFP and Ubi>Kib-Halo localized at the junctional and medial cortex, as observed previously (Figs. 1B & C; Su et al., 2017; Tokamov et al., 2021). Although both Kib and F-actin localize junctionally, Kib’s punctate pattern is distinct from the more uniformly distributed F-actin (Figs. 1B-C’’). In contrast, medial Kib distribution seemed superficially similar to that of F-actin. Indeed, close examination revealed that both Kib::GFP and Ubi-Kib-Halo localized adjacent to or decorated medial F-actin structures (Figs. 1B-C’’), suggesting that Kib might associate with the apical actomyosin network.

We next examined Kib subcellular dynamics with F-actin or myosin via time lapse imaging. Because endogenous Kib is expressed at low levels in wing imaginal discs, we used UASp-Kib-GFP, which produces functional Kib with normal subcellular localization(Tokamov et al., 2021). We observed coordinated movement of Kib with F-actin, and Kib punctae often appeared to be carried by F-actin from the junctional to medial cortex (Fig. 1D, Movie 1).

Similarly, time lapse imaging of Kib-Halo with GFP-tagged Spaghetti squash (myosin regulatory light chain in *Drosophila*, Sqh-GFP) revealed coupled movement of both proteins at the medial cortex (Figs. 1E-F, Movie 2). Together, these results suggest that Kib is associated with the apical actomyosin network and that actomyosin organization might control Kib distribution between the junctional and medial cortex.

### Osmotic shifts lead to coordinated changes in actomyosin and Kib distribution

To further test the relationship between actomyosin dynamics and Kib localization, we sought to acutely alter actomyosin organization and examine any potential changes in Kib localization. Changes in osmolarity have been shown to affect F-actin organization in cultured cells (Cornet et al., 1994; Di Ciano et al., 2002; Pietuch et al., 2013; van Loon et al., 2020), so we decided to use this approach on explanted wing imaginal tissues. In wing imaginal discs incubated under isotonic conditions, both F-actin and myosin are apically enriched, where they are partitioned into junctional and medial pools (Figs. 2A-A’, E-E’). Strikingly, incubating wing imaginal tissues in a hypertonic solution led to medial enrichment of F-actin (Figs. 2B-B’). Ecad localization was unaffected, indicating that this effect was not due to a general collapse of cell-cell junctions (Fig. 2B). Conversely, under hypotonic conditions F-actin accumulated predominantly at the junctional cortex (Figs. 2C-C’). Importantly, the osmotic shifts did not lead to any detectible tissue ruptures and the effects were readily reversible (data not shown). Quantification of F-actin distribution revealed a decrease and increase in junctional/medial F-actin intensity under hypertonic and hypotonic conditions, respectively (Fig. 2D).

**Figure 2:**
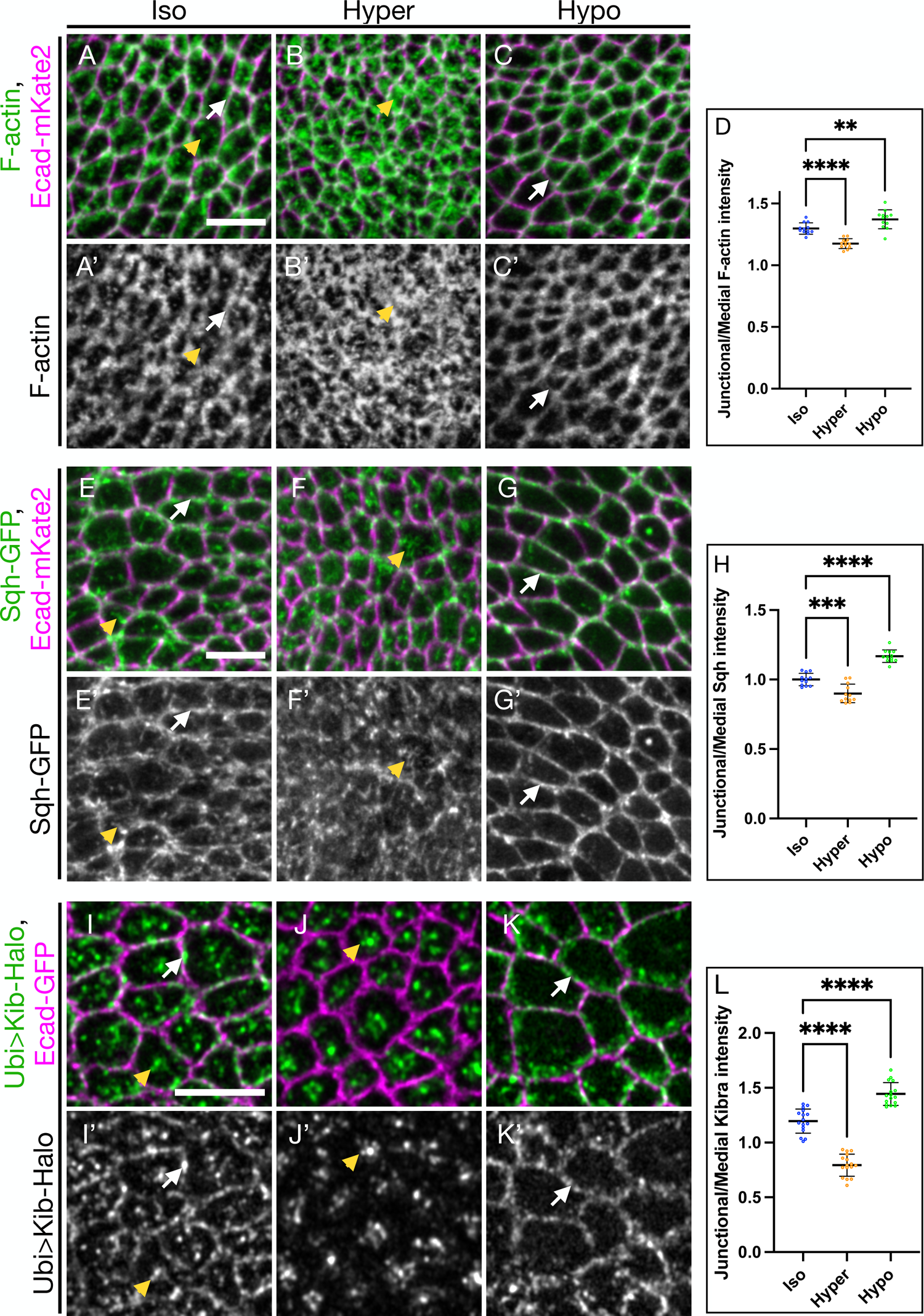
Osmotic shifts lead to coordinated changes in actomyosin and Kib distribution. A-C’) Apical projections of wing imaginal tissues expressing Ecad-mKate2 and an F-actin marker, sGMCA. F-actin is both junctional (white arrows) and medial (yellow arrowhead) under isotonic conditions (A & A’); under hypertonic conditions (B & B’), F-actin accumulates more medially; in contrast, under hypotonic conditions (C-C’), F-actin is more junctional. D) Quantification of junctional/medial F-actin intensity under osmotic shifts. E-G’) Apical projections of wing imaginal tissues expressing Ecad-mKate2 and Sqh-GFP. Sqh is both junctional and medial under isotonic conditions (E & E’); Sqh becomes mostly medial and is decreased at the junctions under hypertonic conditions (F & F’); in contrast, Sqh is more junctional under hypotonic conditions (G-G’). H) Quantification of junctional/medial Sqh intensity under osmotic shifts. I-K’) Under isotonic conditions (I and I’), Kib is both junctional and medial. Similar to F-actin and myosin, Kib becomes predominantly medial under hypertonic conditions (J and J’) and mostly junctional under hypotonic conditions (K and K’). L) Quantification of junctional/medial Kib intensity under osmotic shifts. Scale bars = 5µm. Statistical significance was calculated using One-way ANOVA followed by Tukey’s HSD test.

Similar to the changes in F-actin organization, myosin also became more medially enriched under hypertonic conditions, though it was more prominently lost from the junctions (Figs. 2F-F’). On the other hand, hypotonic shift led to significant junctional accumulation of myosin (Figs. 2G-G’, H). These results show that in the wing imaginal epithelium, changes in extracellular osmolarity can be used to acutely shift subcellular actomyosin distribution between junctional and medial cortical pools.

We next examined whether osmotic shifts affect Kib localization. Compared to the normal junctional and medial Kib distribution under isotonic conditions (Figs. 2I-I’), Kib strongly accumulated medially and became almost undetectable at the junctional cortex under hypertonic conditions (Figs. 2J-J’). In sharp contrast, Kib localized mostly at the junctional cortex under hypotonic conditions (Figs. 2K-K’). Quantification of Kib organization revealed that as in the case of F-actin and myosin, junctional/medial Kib fluorescence decreased under hypertonic and increased under hypotonic conditions relative to isotonic controls (Fig. 2L). Additionally, time lapse imaging of Sqh-GFP with Kib-Halo revealed strong spatial and temporal correlation in Kib and myosin rearrangement upon osmotic shifts. Under the hypotonic shift, myosin and Kib simultaneously localized at the junctional cortex (Movie 3). Conversely, shift to hypertonic medium led to dynamic medial myosin flows that appeared to cluster Kib at the medial cortex (Movie 4). These observations suggest that changes in Kib localization are driven by changes in actomyosin organization.

The drastic changes in Kib localization under osmotic shifts made us wonder if all cortical proteins could be similarly affected by these manipulations. We examined the localization of a spectraplakin Short stop (Shot), which links the minus-ends of apico-basal microtubule arrays to the apical actin cortex in polarized epithelia. Similar to Kib, Shot localizes apically in epithelial cells and can re-localize from the junctional to the medial cortex (Röper and Brown, 2003; Booth et al., 2014). However, in marked contrast to Kib, Shot seemed to enrich more junctionally under hypertonic conditions and accumulated mostly in medial clusters under hypotonic conditions (Figs. S1B-C’). We also examined the localization of Ex and found that unlike Kib, Ex remained predominantly junctional under osmotic shifts (Figs. S1D-D’’). These observations suggest that changes in actomyosin organization could specifically modulate cortical Kib distribution.

### Medial Kib localization is mediated via actomyosin dynamics

Given the strong correlation between actomyosin and Kib distribution under osmotic shifts, we next asked if changes in Kib localization were mediated by actomyosin flows. To this end, we sought to acutely stabilize F-actin using Jasplakinolide (Jasp; Bubb et al., 1994), and examine the effect on Kib localization. Relative to control tissues, F-actin intensity increased dramatically in tissues treated with Jasp, confirming that Jasp potently stablizes F-actin in the wing imaginal discs (Figs. S2A-C). Treating tissues with Jasp under isotonic conditions was sufficient to increase junctional/medial Kib distribution (Figs. 3A-B’, E). Timelapse imaging of sGMCA with Ubi>Kib-Halo showed that medial Kib punctae were dynamic and moved together with F-actin in control tissues, whereas both Kib and F-actin movement was significantly impaired upon treatment with Jasp (Movie 5).

**Figure 3:**
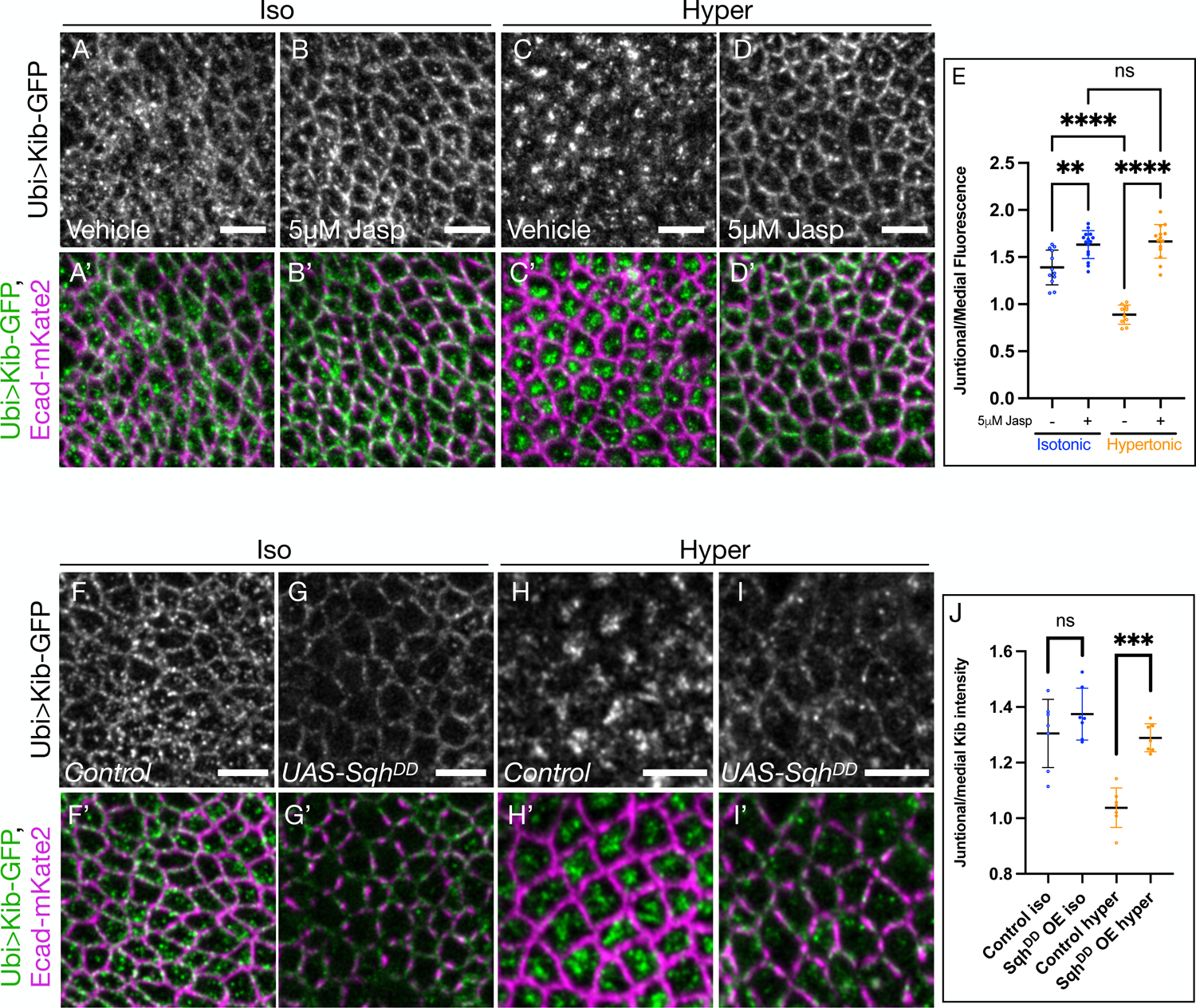
Medial Kib accumulation is mediated via actomyosin dynamics. A-E) Stabilizing F-actin prevents medial Kib localization. Compared to isotonic controls (A and A’), treating wing imaginal tissues with 5µM Jasp results in more junctional Kib (B, B’, E) While Kib is mostly medial under hypertonic conditions (C and C’), treatment with 5µM Jasp blocks the effect of the hypertonic shift on Kib localization (D-E). Scale bars = 5µm. F-J) Dynamic myosin phosphorylation is required for medial Kib accumulation. *UAS-Sqh^DD^* was expressed for 14h using *hh>Gal4* driver and *Gal80^ts^.* Sqh^DD^ expression leads to slightly more junctional Kib under isotonic conditions (F-G’, J) and strongly inhibits medial Kib relocalization under hypertonic conditions (H-J’). Scale bars = 5µm. Statistical significance was calculated using One-way ANOVA followed by Tukey’s HSD test.

We then asked if stabilizing F-actin could prevent medial Kib enrichment induced by the hypertonic shift. Indeed, while Kib accumulated medially in a control hypertonic solution, this relocalization was effectively blocked by the addition of Jasp (Figs. 3C-E). To test if F-actin dynamics were required to maintain the medial Kib pool, we first concentrated Kib medially by incubating tissues in a hypertonic solution and then transferred them into a hypertonic solution with or without Jasp. Treatment with Jasp reversed hypertonically induced medial Kib accumulation (Figs. S2D-E’), suggesting that F-actin dynamics not only accumulate but also maintain Kib at the medial cortex.

We also wondered if inhibiting medial myosin dynamics could have a similar effect on Kib localization. Dynamic phosphorylation-dephosphorylation cycling of the myosin regulatory light chain is required for medial pulsatile myosin activity during morphogenesis(Vasquez et al., 2014). Therefore, we examined Kib localization upon ectopic expression of a phosphomimmetic version of Sqh (Sqh^DD^; Mitonaka et al., 2007). As with F-actin stabilization under Jasp treatment, ectopic Sqh^DD^ expression resulted in significant stabilization of myosin, as evidenced by the increased intensity of GFP-tagged myosin heavy chain (Zip-GFP, Figs. S2F-H). Ectopic Sqh^DD^ expression under isotonic conditions also resulted in a more junctional appearance of Kib (Figs. 3F-G’), though the effect was not quantitatively significant (Fig. 3J), likely because the transgene was expressed transiently in the background of the endogenous Sqh. However, Sqh^DD^ expression significantly blocked medial Kib localization under hypertonic conditions (Figs. 3H-I’, J). Similarly, transient depletion of Sqh blocked medial Kib accumulation both under isotonic and hypertonic conditions (Figs. S2I-J’). Collectively, our data suggest that Kib accumulation at the medial cortex is promoted and actively maintained by medial actomyosin dynamics.

### Actomyosin-driven medial Kib assembles a Hippo signaling complex

Previous work using genetic manipulations has shown that Kib is a key protein that recruits other Hippo pathway components into a signaling complex at the medial cortex, and that medially-localized Kib is more active in restricting growth (Su et al., 2017). However, we wondered if Hippo complex assembly could also occur when Kib is acutely accumulated at the medial cortex upon hypertonic shift. Activation of Hippo signaling by Kib requires the recruitment of Mer, and Mer helps recruit the core kinase cassette to the medial cortex (Su et al., 2017). Under isotonic conditions, we observed some colocalization between Kib and Mer at the medial and junctional cortex (Figs. 4A-A’’). Under hypertonic conditions, both proteins strongly accumulated at the medial cortex and their colocalization was more pronounced (Figs. 4B-B’’), suggesting that actomyosin-driven medial Kib can rapidly recruit Mer into a signaling complex. Conversely, under hypotonic conditions Kib and Mer were both at the junctional cortex where their colocalization was less apparent (Figs.4C-C’’).

**Figure 4:**
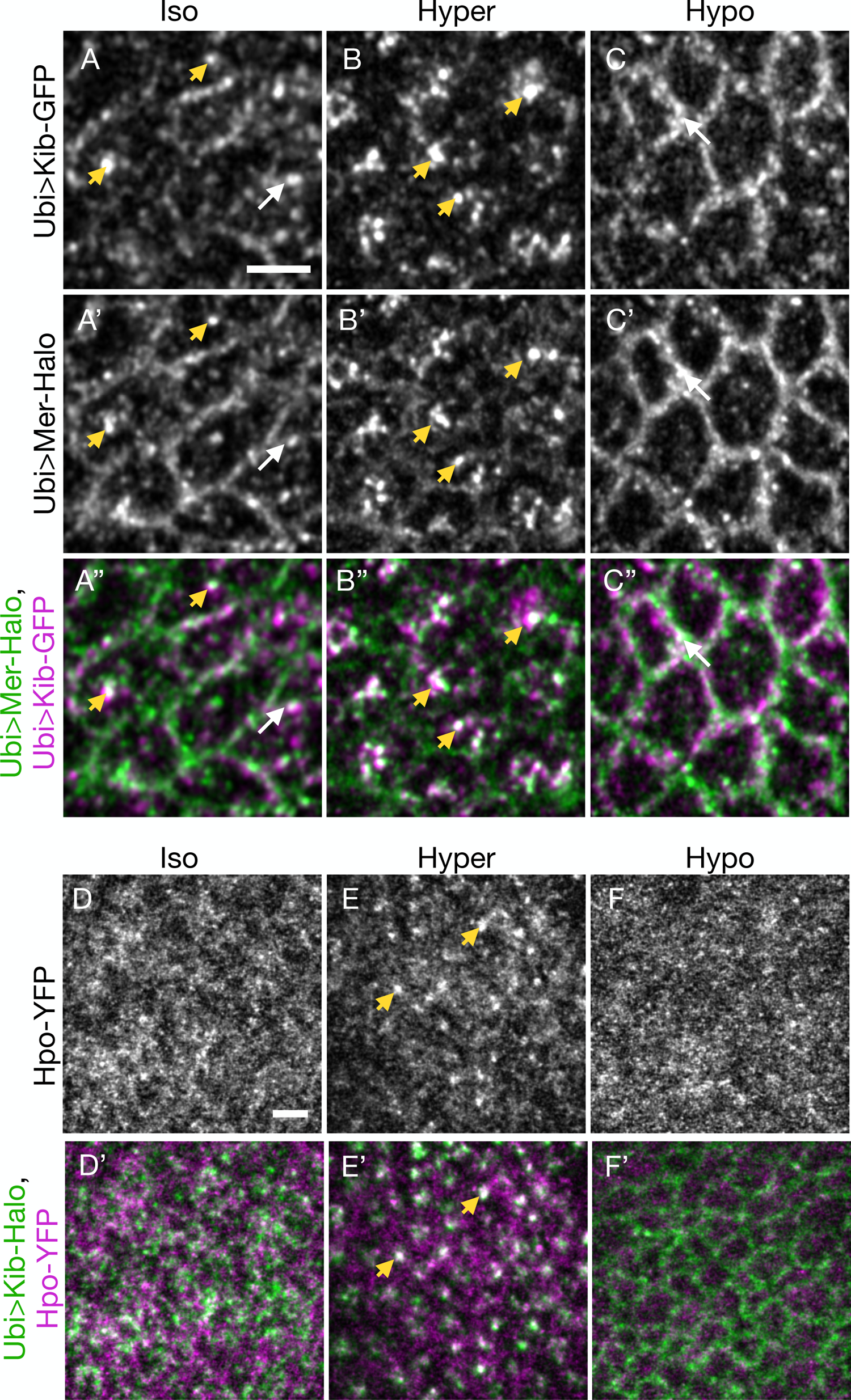
Actomyosin-driven medial Kib assembles a Hippo signaling complex. A-C’’) Mer displays some co-localization with Kib at the junctional (white arrows) and medial (yellow arrowheads) cortex under isotonic conditions (A-A’’). Under hypertonic conditions, Mer is strongly recruited to medial foci with Kib (B-B’’). Conversely, under hypotonic conditions, both Mer and Kib localize mainly at the junctional cortex (C-C’’). D-F’) Hpo is normally diffuse under isotonic conditions (D-D’). However, under the hypertonic shift, Hpo is recruited to the medial cortex with Kib (E-E’). In contrast, Hpo remains diffuse under hypotonic conditions (F-F’). Scale bars = 3µm.

Next, we examined the effect of Kib relocalization under osmotic shifts on Hpo recruitment using endogenously expressed Hpo-YFP. Under isotonic conditions, Hpo-YFP appeared mostly diffuse, with no discernable localization pattern (Figs. 4D-D’). However, as was observed with Mer, distinct Hpo foci formed and colocalized with medial Kib upon hypertonic shift (Figs. 4E-E’). Conversely, shift to hypotonic conditions led to predominantly diffuse Hpo-YFP, with no detectible junctional localization (Figs. 4F-F’). Collectively, these data suggest that acute medial Kib accumulation via actomyosin flows promotes Hippo complex assembly.

### Crb controls Kib localization via actomyosin organization and cortical tethering

To this point, our results demonstrate that actomyosin flows promote medial Kib accumulation and Hippo complex formation. However, our observations that Kib accumulates junctionally when actomyosin dynamics are inhibited suggests that Kib could be tethered at the junctional cortex. We previously showed that Crb is required for junctional Kib localization (Su et al., 2017), prompting us to further investigate the relationship between Crb and Kib.

Since loss of Crb results in similar medial Kib accumulation as observed under the hypertonic shift (Fig. S3A-B’), we wondered if the actomyosin network was also altered in cells lacking Crb. Previous studies reported that loss of Crb leads to increased medial actomyosin organization (Flores-Benitez and Knust, 2015; Salis et al., 2017). Consistent with those reports, we found that F-actin organization in *crb*-null clones was significantly more medial (Figs. S3C-C’). We also found that similar to hypertonic conditions, loss of Crb disrupted junctional myosin organization and led to increased medial myosin accumulation (Figs. S3D-D’, Movie 6). These observations suggest that medial accumulation of Kib in cells lacking Crb could be caused by increased medial actomyosin flows. Indeed, stabilizing F-actin via Jasp treatment blocked medial Kib accumulation in *crb-*depleted cells (Figs. S3E-F’). Interestingly, while Jasp treatment resulted in more junctional Kib localization in control cells, it failed to restore junctional Kib in cells depleted of Crb (Figs. S3E-F’). These results suggest that Crb can regulate Kib via two simultaneous mechanisms: 1) by inhibiting medial actomyosin flows and 2) by providing a tether to maintain Kib at the junctional cortex.

How could Crb tether Kib junctionally? Crb has a short intracellular region containing only two known functional motifs. The FERM-binding motif, which interacts with Ex, Moesin, and Yurt (Tepass, 2012), is not required for Kib junctional localization (Su et al., 2017). Consistent with this, while Crb depletion led to more medial Kib, loss of Ex had no effect on Kib localization (Figs. S4A-B’’). On the other hand, Crb’s PDZ-binding motif interacts with Sdt/Patj and aPKC/Par6 complexes, which in turn could recruit Kib since Kib itself does not contain a PDZ domain. Interestingly, loss of Sdt or Patj had no effect on Kib localization (Figs. S4C-D’’), suggesting that Crb could tether Kib at the junctional cortex via aPKC/Par6.

### aPKC tethers Kib at the cell cortex in cultured cells

Several lines of evidence suggest that aPKC could tether Kib cortically downstream of Crb. First, Kib contains an aPKC binding motif and is known to physically interact with aPKC in *Drosophila* and mammalian cells (Büther et al., 2004; Yoshihama et al., 2011; Jin et al., 2015). Second, aPKC is highly enriched at the junctional cortex, where it partially colocalizes with Kib punctae (Figs. S4E-E’’). Third, we found that in the wing imaginal disc, loss of Crb leads to a significant decrease in cortical aPKC (Figs. S4F-F’’), consistent with previous observations and reports that Crb forms a complex with Par6 and aPKC and promotes aPKC activity (Genevet et al., 2009; Morais-de-Sá et al., 2010; Walther and Pichaud, 2010; Dong et al., 2020).

To test the tethering role of aPKC, we first used cultured Schneider’s 2 (S2) cells to ask if aPKC can recruit Kib to the cell cortex. When expressed by itself, Kib normally aggregates in cytosolic foci and almost never appears at the cortex (Baumgartner et al., 2010; Figs. 5A & E). In contrast, co-expression of Par6 and aPKC, both of which localize cortically in S2 cells, led to recruitment of Kib to the cell cortex in ∼40% of cells, as evidenced by the appearance of a bright rim around the cell optically sectioned through the equatorial plane (Figs. 5B-B’’, E). To ask if Crb can modify cortical Kib recruitment via Par6/aPKC, we expressed the intracellular region of Crb (Crb^i^) alone or together with Par6 and aPKC and examined Kib localization. Crb^i^ by itself did not recruit Kib to the cortex (Figs. 5C-C’, E). However, Kib localized cortically in over 60% of cells when Crb^i^, Par6, and aPKC were co-expressed (Figs. 5D-D’’, E).

**Figure 5.**
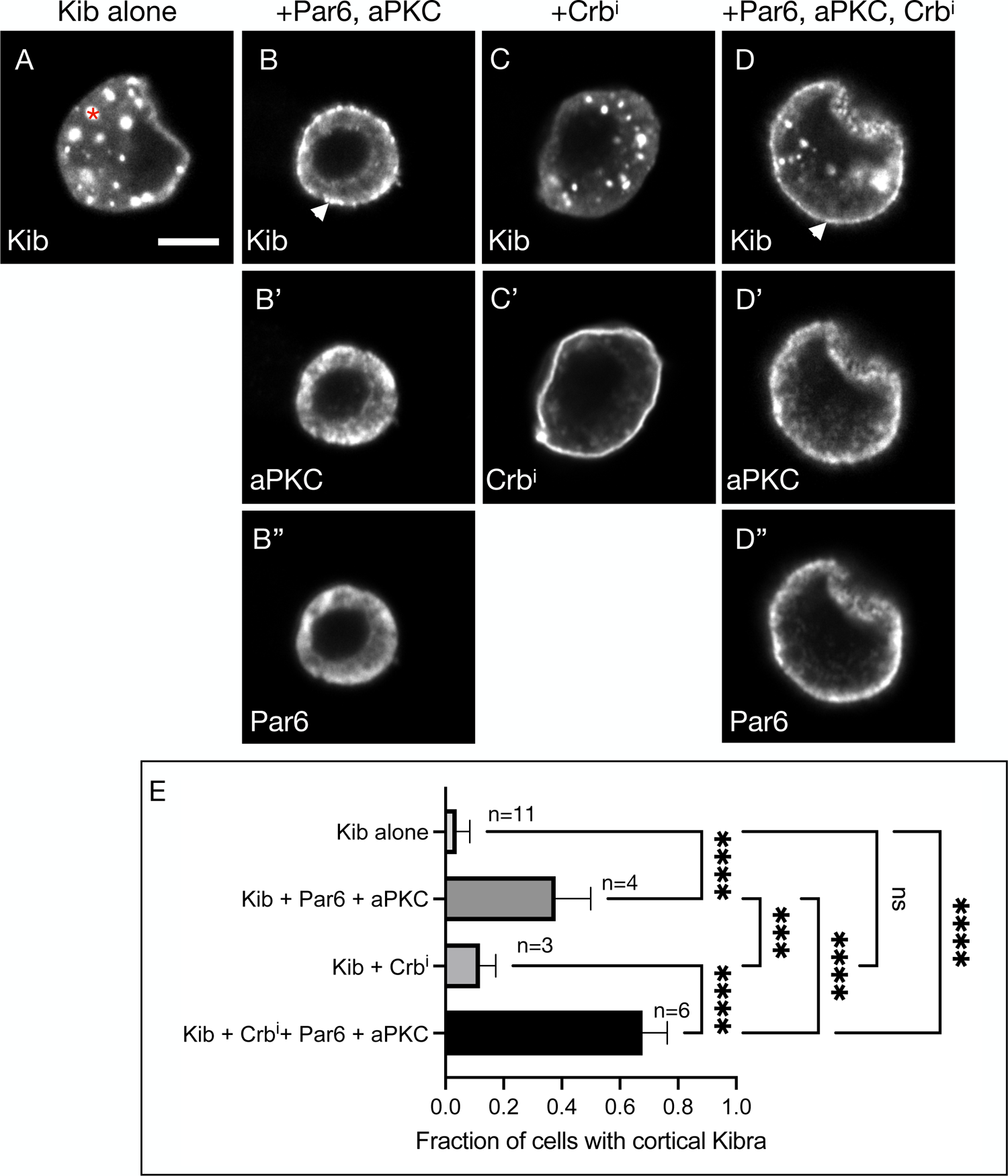
aPKC tethers Kib at the cell cortex in S2 cells. A) Expressed by itself, Kib often aggregates in cytoplasmic foci (asterisk) in cultured S2 cells. Scale bar = 5µm. B-B’’) Co-expression of aPKC and Par6 leads to cortical Kib recruitment (arrowhead in B). C-C’) Expression of Crb^i^ alone does not result in cortical Kib recruitment. D-D’’) Addition of Crb^i^ enhances Par6/aPKC-mediated cortical Kib recruitment. E) Quantification of the fraction of cells displaying cortical Kib under the conditions shown in A-D’’. Statistical significance was calculated using One-way ANOVA followed by Tukey’s HSD test.

If Crb functions to stabilize aPKC, then cortically anchoring aPKC by other means should also lead to robust cortical Kib recruitment. To test this idea, we adapted a previously published method of inducing polarity in S2 cells that expresses aPKC fused to the cytoplasmic terminus of the homophilic adhesion protein, Echinoid (Ed, Johnston et al., 2009). Expressing this fusion construct (Ed:GFP:aPKC) in S2 cells leads to cell clustering (via the Ed extracellular domain) and enrichment of aPKC at cell-cell contacts (Fig. 6A). As a control, we used an analogous construct lacking aPKC (Ed:GFP). Using this approach in combination with Kib truncations, we asked if Ed:GFP:aPKC could recruit Kib to cell-cell contacts and if so, whether the aPKC-binding region of Kib is necessary and sufficient for this interaction (Fig. 6A-B). As expected, wild-type Kib (Kib^WT^) appeared largely cytoplasmic in cells clustered via Ed:GFP, indicating that formation of cell-cell contacts per se does not lead to cortical Kib localization (Figs. 6C-C’’, G). In sharp contrast, Kib was recruited to cell-cell adhesion sites upon Ed:GFP:aPKC expression (Figs. 6D-D’’, G). Furthermore, a Kib variant lacking the aPKC-binding region (Kib^!aPKC^) failed to localize at the cell-cell contacts (Figs. 6E-E’’, G), whereas a C-terminal fragment of Kib containing the aPKC-binding region (Kib^858-1288^) was robustly recruited by Ed:GFP:aPKC (Figs. 6F-F’’, G). Together, these results suggest that aPKC recruits Kib to the cell cortex and that the aPKC-binding motif of Kib is both necessary and sufficient for this interaction.

**Figure 6.**
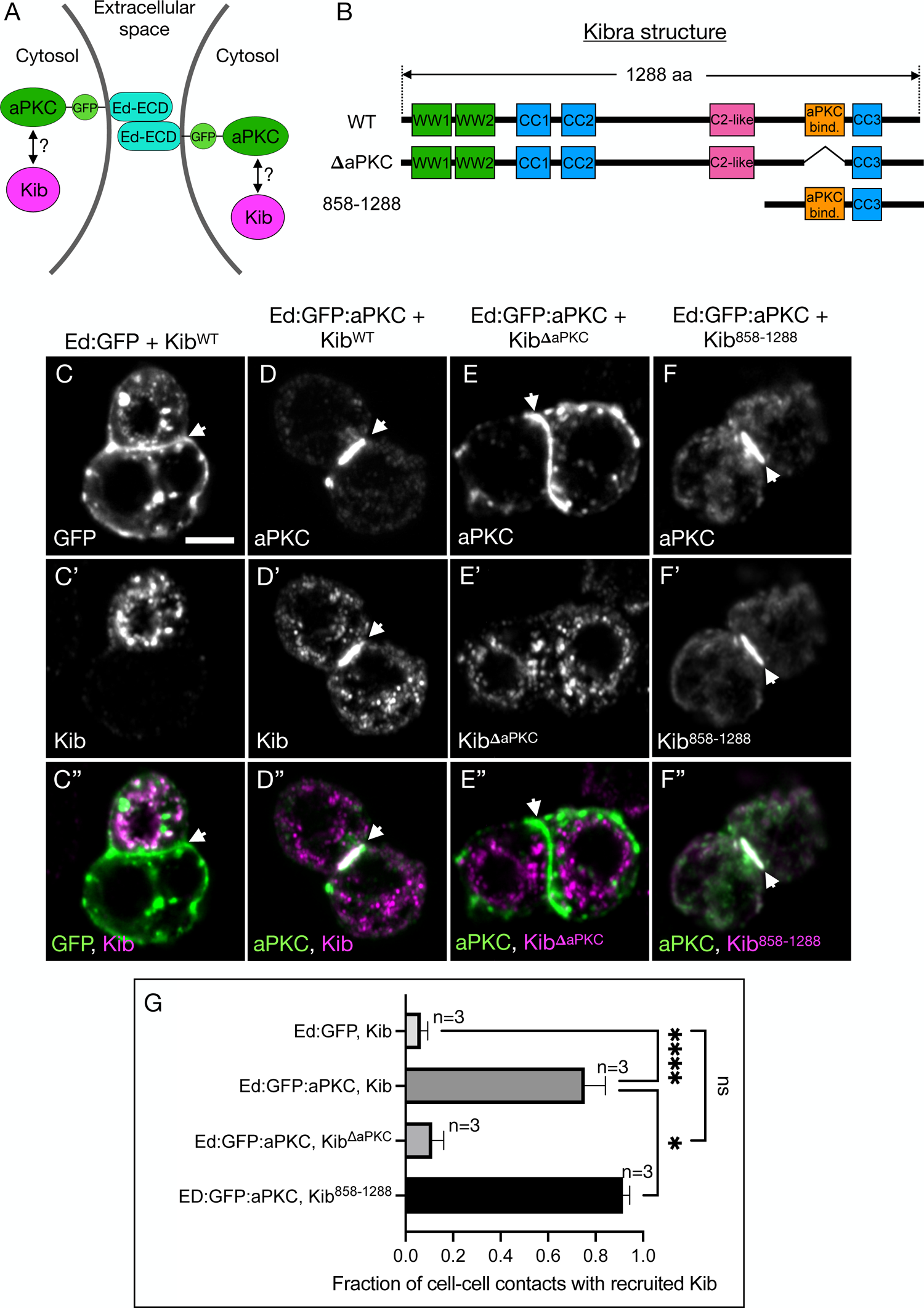
The aPKC-binding domain in Kib is necessary and sufficient for its cortical recruitment by aPKC. A) A schematic of two cells clustered via Ed extracellular domain (Ed-ECD) fused to aPKC on the intracellular side. B) Protein structure cartoons of wild type (WT) Kib, Kib lacking the aPKC-binding motif (ΔaPKC), or Kib containing the last 431 amino acids (858-1288). C-C’’) In control cell clusters generated via Ed:GFP, Kib aggregates predominantly in cytoplasmic foci and is absent from cell-cell contacts (arrowhead). Scale bar = 5µm. D-D’’) When cells are clustered via Ed:GFP:aPKC, Kib is recruited to cell-cell contacts. E-E’’) Ed:GFP:aPKC fails to recruit Kib^ΔaPKC^ to cell-cell contacts. F-F’’) Ed:GFP:aPKC robustly recruits Kib^858-1288^ to cell-cell contacts. G) Quantification of the fraction of cell-cell contacts with recruited Kib under the conditions shown in C-F’’. Statistical significance was calculated using One-way ANOVA followed by Tukey’s HSD test.

### aPKC tethers Kib at the junctional cortex to limit Kib-mediated Hippo signaling

We next tested if aPKC tethers Kib at the junctional cortex in the wing imaginal discs and the potential functional significance of this interaction. Loss of aPKC activity disrupts cell polarity and constitutive depletion of aPKC in the wing imaginal discs leads to cell death (Sotillos et al., 2004), making it difficult to assess the role of aPKC in localizing Kib using genetic manipulations. Therefore, we used a previously published analog-sensitive allele of aPKC, *aPKC^as4^*, which produces a functional kinase that can be acutely inhibited by a small molecule 1NA-PP1 (Hannaford et al., 2019). Treatment of wing imaginal discs homozygous for *aPKC^as4^*with 1NA-PP1 for only 15 min severely disrupted normal cortical aPKC localization (Figs. S5A-B), similar to what was previously observed in Drosophila larval neuroblasts (Hannaford et al., 2019). Importantly, 1NA-PP1 had no effect on aPKC or Kib-GFP distribution in the background of wild-type *aPKC* (Fig. S5C-E’), indicating that the effect of 1NA-PP1 is specific to *aPKC^as4^*.

To test the effect of aPKC inhibition on Kib localization, we treated wing imaginal explants expressing Ubi>Kib-GFP in *aPKC^as4^* background with 1NA-PP1 for 15 min. Strikingly, this resulted in Kib accumulation almost exclusively at the medial cortex (Figs. 7A-B’), supporting the role of aPKC as a junctional tether for Kib. Importantly, Mer also accumulated medially under 1NA-PP1 treatment whereas Ex remained junctional (Figs. S6A-D’), suggesting that aPKC inhibition affects Kib-mediated signaling specifically and consistent with the previous observation that acute aPKC inhibition does not affect apical Crb localization (Osswald et al., 2022).

**Figure 7.**
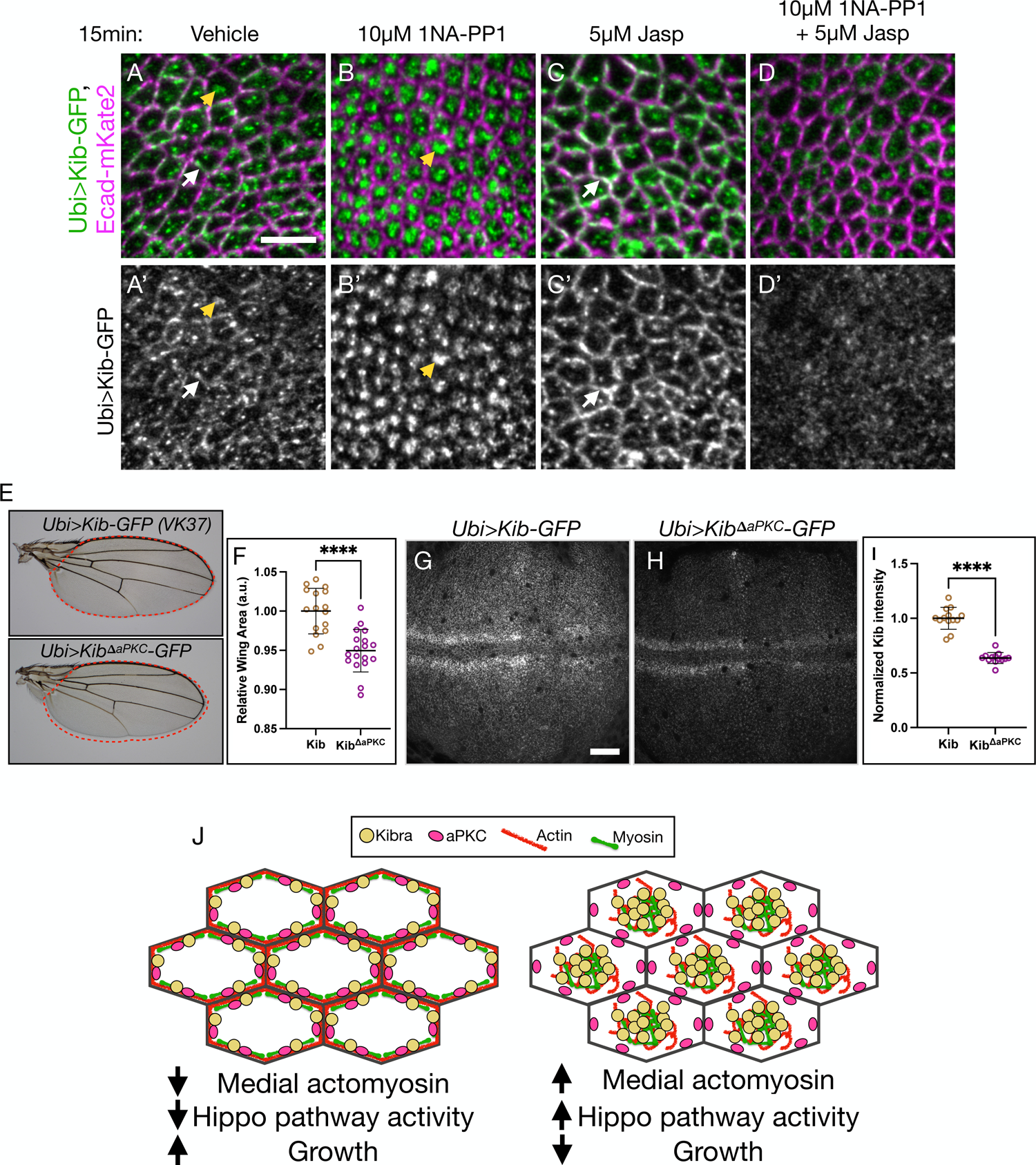
Tethering by aPKC limits Kib-mediated Hippo pathway activation. A-D’) In tissues homozygous for *aPKC^as4^*allele, Kib localizes at the junctional (white arrows) and medial (yellow arrowheads) cortex under control conditions (A-A’) but is predominantly medial under aPKC inhibition with 1NA-PP1 (B-B’). Treatment with Jasp leads to more junctional Kib accumulation (C-C’). Under simultaneous treatment with 1NA-PP1 and Jasp, Kib fails to accumulate medially or junctionally (D-D’). Scale bar = 5µm. E-F) Wings ectopically expressing Kib^ΔaPKC^ (*Ubi>Kib^ΔaPKC^-GFP*) are slightly smaller the ones expressing wild-type Kib (*Ubi>Kib-GFP*). G-I) Wild-type Kib (G) is more stable than Kib^ΔaPKC^ (H). Scale bar = 20µm. Transgenes in E-H are identically expressed. Statistical significance in F & I was calculated using Mann-Whitney test. J) A cartoon model of Kib regulation via apical polarity and actomyosin flows. Under low medial actomyosin flows, aPKC tethers and inhibits Kib at the junctional cortex, resulting in more growth. Stronger medial actomyosin flows untether Kib from the junctional cortex and accumulate it medially, where it recruits associated components to promote Hippo signaling and inhibit growth.

As with loss of Crb, aPKC inhibition affected actomyosin organization. Recent studies in *Drosophila* reported that acute aPKC inactivation leads to increased medial myosin activity and apical constriction in embryonic and the follicular epithelial cells (Hannaford et al., 2019; Biehler et al., 2021; Osswald et al., 2022). In agreement with these reports, we observed increased medial myosin and F-actin accumulation in the wing imaginal disc cells under aPKC inhibition (Figs. S6E-H’), suggesting that changes in actomyosin organization could be responsible for medial Kib accumulation under aPKC inhibition. Consistent with this hypothesis, inhibition of aPKC in the presence of Jasp resulted in diffuse Kib that failed to accumulate at the junctional or medial cortex (Figs. 7C-D’). Collectively, these results strongly suggest that Kib cortical distribution is controlled by the opposing action of aPKC, which tethers Kib at the junctional cortex, and actomyosin flows, which accumulate Kib medially.

What could be the potential function of junctional Kib recruitment by aPKC? Previous work suggested that loss of Crb enhances Kib’s ability to activate the Hippo pathway and suppress tissue growth (Su et al., 2017). These observations suggest that aPKC could inhibit Kib via junctional tethering. To test this idea, we compared adult wing size from flies expressing either wild-type Kib or Kib^ΔaPKC^ under the ubiquitin promoter and from identical genomic locations. Flies expressing Kib^ΔaPKC^ had mildly undergrown wings compared to those expressing wild-type Kib (Figs. 7E-F). Additionally, despite its heightened ability to suppress growth, Kib^ΔaPKC^ abundance was significantly lower than that of wild-type Kib (Figs. 7G-I), suggesting that Kib^ΔaPKC^ is intrinsically more active than wild-type Kib. We observed similar differences in adult wing growth and protein abundance when wild-type Kib and Kib^ΔaPKC^ were expressed under UAS control using a wing-specific driver *Nubbin>Gal4* (Figs. S5F-J), indicating that these differences were not due to the ubiquitous expression of the transgenes. Together, these results suggest that aPKC-mediated tethering of Kib at the junctional cortex limits Kib-mediated Hippo pathway activation.

## Discussion

Proper spatial organization of subcellular signaling components is critical for controlling the specificity and amplitude of signaling pathways (Corbit et al., 2005; Lancaster et al., 2011; Rys et al., 2015). Despite our long-standing knowledge that upstream Hippo pathway regulators are enriched at the apical cortex of epithelial cells, our understanding of how these proteins are organized and the importance of apical distribution in regulating their signaling output is far from complete. In this study, we focused on the cortical distribution of Kib, a key upstream Hippo pathway activator that is organized not only along the apico-basal axis, with strong enrichment apically, but also across the apical cortex into junctional and medial pools. We demonstrate that such distribution of Kib is achieved by opposing influences from the apical polarity and actomyosin networks. While the apical polarity network, at least in part via aPKC, tethers Kib at the junctional cortex to silence its activity, centripetal actomyosin flows promote Hippo signaling via accumulation of Kib and the associated Hippo signaling components at the medial cortex (Fig. 7J).

Our study identifies a previously unrecognized role of the medial actomyosin network in regulating the Hippo pathway. Actomyosin-generated cortical tension is a conserved stimulus that inhibits Hippo signaling and promotes Yki activity and growth. However, while cortical tension inhibits Hippo signaling at the kinase level via Ajuba-mediated sequestration of Wts (Rauskolb et al., 2014; Ibar et al., 2018), how the actomyosin cytoskeleton might affect upstream Hippo pathway regulators is unknown. A striking observation in our study is that manipulations that affect actomyosin organization and cortical tension can drive dramatic changes in Kib localization (Figs. 2-4, Movie 3, Movie 4). Given that myosin organization and dynamics are known to be influenced by cortical tension (Fernandez-Gonzalez et al., 2009; Duda et al., 2019), we propose that cortical tension could regulate medial Kib localization via modulation of actomyosin flows. Specifically, we propose that higher cortical tension leads to junctional actomyosin enrichment and decreased medial flows, resulting in less medial Kib and attenuated Hippo signaling (Fig. 7J). The opposite occurs under lower cortical tension. This mechanism could be important in the developing wing imaginal disc, where higher tension at the tissue periphery is thought to inhibit the Hippo pathway to promote growth via Yki (Aegerter-Wilmsen et al., 2007, 2012; LeGoff et al., 2013; Mao et al., 2013; Hariharan, 2015; Pan et al., 2016). It could also function more generally during tissue morphogenesis driven by actomyosin dynamics, such as during the apical constriction that drives furrow formation in a range of epithelia. Tension-driven actomyosin flows have also been reported to transport ZO-1 clusters to tight junctions during zebrafish epiboly (Schwayer et al., 2019), suggesting that this could be a general mechanism to control cortical protein distribution during tissue growth and morphogenesis.

How could actomyosin-mediated medial Kib accumulation promote Kib activity? We observed that actomyosin flows drive translocation of Kib punctae from the junctional cortex, where aPKC is enriched, to the medial cortex, where Kib punctae appear to form larger assemblies (Movies 1 & 4). Furthermore, the formation of such medial assemblies is enhanced by treatment with hypertonic medium, Crb depletion, and acute aPKC inhibition, all of which also increase medial actomyosin organization, and inhibited when the actomyosin network is stabilized (Figs. 2, 3, S3, & 7). We propose that medial actomyosin flows promote Hippo signaling by driving Kib coalescence and Hippo signaling complex formation at the medial cortex, away from Kib’s junctional inhibitors, including aPKC (Fig. 7J). A recent study also argued that medial Kib (but not junctional) undergoes phase separation, thereby promoting its ability to assemble pathway components (Wang et al., 2022). Our results are consistent with this possibility, though they do not directly confirm it. The kinetics of Kib cluster formation and its regulation by actomyosin flows will have to be resolved by further studies.

A limitation in this study (and the field at large) is the lack of a biosensor that directly reports Kib-mediated Hippo pathway activation in living cells. However, several lines of evidence suggest that medial accumulation promotes Kib activity. First, loss of Crb, which results in predominantly medial Kib, enhances Kib-mediated signaling (Su et al., 2017). Second, the size and dynamics of medial Kib assemblies are similar to biomolecular condensates formed by Kib in mammalian cells, and phase transition was suggested to promote Kib activity at the medial cortex (Wang et al., 2022). Third, Hpo co-accumulates with Kib at the medial cortex but not at the junctional cortex (Fig. 4; Su et al., 2017). Because Hpo dimerizes and trans-autoactivates (Jin et al., 2012), it is likely that medial Kib accumulation provides the means of increasing Hpo activation via local concentration, as has been suggested for their mammalian counterparts (Wang et al., 2022). Fourth, Merlin, which promotes Kib’s ability to assemble pathway components, also accumulates with Kib medially (Fig. 4; Su et al., 2017). Additionally, given the evidence that medial Kib assemblies are significantly more stable than the junctional Kib pool (Wang et al., 2022), it is possible that medial accumulation promotes Hippo signaling by increasing the cortical dwell time of Kib complexes (Case et al., 2019; Huang et al., 2019).

One of the surprising aspects of our findings is the specificity of actomyosin-mediated regulation of Kib localization. Ex and Kib are similar in size (∼154kDa and 144kDa, respectively), assemble similar signaling complexes, colocalize junctionally, and are known to physically interact in cultured S2 cells (Yu et al., 2010; Genevet et al., 2010; Sun et al., 2015; Su et al., 2017). However, while Kib rapidly relocalizes from the junctional to the medial cortex under hypertonic conditions or aPKC inhibition, Ex remains junctional (Figs. 2, 7, S1D-D’’, S6C-D’). This difference in behavior cannot be explained solely by the difference in tethering strengths because loss of Crb untethers both Ex and Kib from the junctional cortex, but whereas Ex becomes cytoplasmically diffuse, Kib accumulates medially (Robinson et al., 2010; Su et al., 2017, Fig. S3A-B’). These observations are consistent with previous reports indicating that Ex and Kib function in parallel to form separate signaling modules (Baumgartner et al., 2010; Yu et al., 2010; Su et al., 2017; Tokamov et al., 2021), and they also raise the question of how Kib localization could be specifically regulated by the actomyosin flows. Kib lacks an actin-binding domain and our observations of Kib localization with respect to F-actin and myosin suggest that Kib is unlikely to associate tightly with the actomyosin network (Fig. 1). Three non-mutually exclusive possibilities could explain why actomyosin flows accumulate Kib but not Ex at the medial cortex. First, the multivalency of Kib could enable weak interactions with the actomyosin cytoskeleton and/or its regulators, thereby facilitating advective transport by actomyosin flows, as has been shown for Par3 in the *C. elegans* zygote (Munro et al., 2004; Goehring et al., 2011). Second, the interaction between Kib and aPKC could be modulated by factors that also regulate actomyosin dynamics. Third, in addition to its junctional tethering, Kib could be stabilized by a yet unknown anchor at the medial cortex. The tools and framework developed in this study provide a strong foundation for future investigation of these possibilities.

This study highlights the relationship between Kib and aPKC. Our results suggest that junctional tethering by aPKC attenuates Kib’s ability to activate the Hippo pathway. We show that aPKC recruits Kib to the cortex in S2 cells (Figs. 5 & 6), and Kib lacking the ability to bind aPKC is intrinsically more active in vivo (Fig. 7). Because Crb is necessary for aPKC recruitment to the junctional cortex, these observations are consistent with previous data showing that loss of Crb potentiates Kib-mediated Hippo pathway activation (Su et al., 2017). Similar to our observations of the relationship between Kibra and aPKC in the wing, previous work in the *Drosophila* fat body suggested that aPKC inhibits Kib’s ability to promote starvation-induced autophagy (Jin et al., 2015), although it is unclear if Kib-mediated regulation of autophagy involves Hippo signaling. Nonetheless, multiple lines of evidence suggest that aPKC regulates Kibra function through direct interaction.

Our results also suggest that aPKC likely is not the only component that tethers Kib because ectopically expressed Kib^ΔaPKC^ is still able to localize junctionally (Fig. S5K-L). However, the additional Kib tether must, catalytically and/or structurally, depend on aPKC, because simultaneous inhibition of both aPKC and actomyosin dynamics prevents junctional accumulation of full-length Kib normally observed when actomyosin dynamics alone are perturbed. Both Par6 and the *Drosophila* homolog of Par3, Bazooka (Baz), mislocalize under acute aPKC inhibition (Aguilar-Aragon et al., 2018; Hannaford et al., 2019), suggesting that they could assist aPKC in stabilizing Kib at the junctional cortex. On one hand, the involvement of Baz in aPKC-mediated Kib regulation seems unlikely since aPKC was shown to exclude Baz from the marginal zone to the adherens junctions in the follicular epithelium (Morais-de-Sá et al., 2010). On the other hand, this exclusion requires aPKC phosphorylation of Baz on S980, as non-phosphorylated Baz (S980A) associates with Par6/aPKC at the apical membrane (Benton and St Johnston, 2003; Morais-de-Sá et al., 2010). Given that Kib is believed to bind to aPKC’s kinase domain and inhibit aPKC’s kinase activity (Yoshihama et al., 2011, 2013), it is unlikely that aPKC bound to Kib would be able to exclude Par3 from binding Par6. In addition, mammalian Kib also directly interacts with Patj (Duning et al., 2008). Although we find that depletion of Patj has no effect on junctional Kib localization, the existence of multiple interactions stabilizing Kib could obscure the role of individual components, especially under long-term genetic perturbations. Regardless of the molecular details, however, the existence of multiple tethers also suggests that Kib could be associated with a broader network of polarity organizers.

Finally, our work highlights the importance of understanding the role and regulation of Hippo signaling in a broader developmental context. Proper organ development involves robust control of both tissue size and shape, suggesting that these processes must be regulated in a concerted fashion. Cell polarity and actomyosin networks play central roles in the regulation of growth and morphogenesis (Tepass, 2012; LeGoff and Lecuit, 2016; Irvine and Shraiman, 2017; Fomicheva et al., 2020), but our understanding of how the interaction between epithelial architecture and mechanical forces is biochemically translated into intracellular signaling events is currently limited. Our findings that Kib subcellular organization and activity are regulated by the tug of war between apical polarity and actomyosin networks suggest that Kib could serve as a nexus in coordinating growth and morphogenesis. Further investigation of how Kib and other Hippo signaling components fit into the molecular network governing epithelial cell organization will provide valuable insight into the broader functions of the Hippo pathway in development.

## Methods

### Drosophila husbandry

*Drosophila melanogaster* was cultured using standard techniques at 25°C (unless otherwise noted). For *Gal80^ts^* experiments, crosses were maintained at 18°C and moved to 29°C for the duration specified in each experiment.

### Osmotic and pharmacological treatment of wing imaginal tissue explants

All osmotic solutions were prepared fresh per experiment. Schneider’s Drosophila Medium (Sigma) supplemented with 10% Fetal Bovine Serum (Thermo Fisher Scientific) was used as isotonic medium (∼360mOsm). To make a hypertonic solution, the osmolarity of the isotonic solution was increased to ∼460mOsm using 1M NaCl. To make a hypotonic solution, the isotonic medium was diluted with deionized water to 216mOsm.

To inhibit F-actin dynamics, tissues were incubated in an isotonic or hypertonic solution with DMSO or 5µM Jasplakinolide before mounting. To inhibit aPKC, tissues homozygous for *aPKCas4* allele were incubated in an isotonic solution with DMSO or 10µM 1NA-PP1 before mounting. Unless indicated otherwise, all incubations were done for 15 min in a humid chamber, and tissues were imaged live immediately after incubation.

### Live imaging

Except for Fig. S4F-F’’, tissues were imaged live. Live imaging of the wing imaginal discs was adapted from Restrepo et al. (2016). Briefly, wing imaginal tissues were first dissected in Schneider’s Drosophila Medium (Sigma) supplemented with 10% Fetal Bovine Serum (Thermo Fisher Scientific) on a siliconized glass slide. Tissues were then transferred with a pipette in 5-10µl of medium to a glass bottom microwell dish (MatTek, 35mm petri dish, 14mm microwell) with No. 1.5 coverglass. The discs were oriented so that the apical side of the disc proper faced the coverglass. A Millicell culture insert (Sigma, 12mm diameter, 8µm membrane pore size) was prepared in advance by cutting off the bottom legs with a razor blade and removing any excess membrane material around the rim of the insert. The insert was carefully placed into the 14mm microwell space, directly on top of the drop containing the tissues. The space between the insert and the microwell was sealed with ∼15µl of mineral oil and 200µl of isotonic medium was added into the insert chamber. For osmotic shift movies with Sqh-GFP and Ubi>Kib-Halo, isotonic medium was removed and 200µl of media with indicated osmolarity was added into the chamber of the insert immediately before scanning.

An inverted Zeiss LSM880 laser scanning confocal microscope equipped with a GaAsP spectral detector and the Airyscan module was used for all imaging. All timelapse images and still images in Figs. 1, 2I-K’, S1D-D’’, 4, S4E-E’’, and S6E-H’ were taken using the Airyscan in superresolution mode. Timelapse acquisition was performed using the Zeiss Z-Piezo. Images were processed in ImageJ.

### Image analysis and quantification

ImageJ was used for basic image processing, including generation of maximum projections, background subtraction, and Gaussian blur application. To quantify junctional/medial intensity, maximum projections were segmented via Cellpose (using Ecad-mKate2 as the junctional marker) and a standard Scikit watershed algorithm (van der Walt et al., 2014; Stringer et al., 2021). The junctional mask generated by segmentation was then applied to quantify mean junctional Kib intensity, while remaining non-junctional signal was classified as medial. The ratio of mean junctional to medial intensity was then calculated. Plots and statistical analyses of mean fluorescence intensities were generated using GraphPad Prism software.

### Immunostaining of imaginal tissues

In Fig. S4F-F’’, the wing imaginal discs from wandering late third instar larvae were fixed in 2% paraformaldehyde (PFA)/PBS and stained as previously described (McCartney and Fehon, 1996). Primary antibodies and concentrations are listed in Table 1. Secondary antibodies (diluted 1:1000) were from Jackson ImmunoResearch Laboratories.

**Table 1:** Reagents.

| Reagent type (species) or resource | Designation | Source or reference | Identifiers | Additional information |
| --- | --- | --- | --- | --- |
| genetic reagent (*D. melanogaster*) | *Kib::GFP* | (Su et al., 2017) |  |  |
| genetic reagent (*D. melanogaster*) | *UAS-LifeAct-Halo2* | Bloomington Drosophila Stock Center | BL67625 |  |
| genetic reagent (*D. melanogaster*) | *Ubi>Kib-Halo* | This study |  |  |
| genetic reagent (*D. melanogaster*) | *sGMCA* | (Kiehart et al., 2000) |  |  |
| genetic reagent (*D. melanogaster*) | *Sqh-GFP* | Bloomington Drosophila Stock Center | BL57145 |  |
| genetic reagent (*D. melanogaster*) | *Ubi>Kib-GFP* | (Tokamov et al., 2021) |  |  |
| genetic reagent (*D. melanogaster*) | *Ubi>Kib<sup>ΔaPKC</sup>-GFP* | This Study |  |  |
| genetic reagent (*D. melanogaster*) | *Zip-GFP* | (Rauskolb et al., 2014) |  |  |
| genetic reagent<br>(*D. melanogaster*) | *UAS-Sqh<sup>DD</sup>* | (Mitonaka et al., 2007) |  |  |
| genetic reagent<br>(*D. melanogaster*) | *Ecad-3XmKate2* | (Pinheiro et al., 2017) |  |  |
| genetic reagent<br>(*D. melanogaster*) | *Ex-YFP* | (Su et al., 2017) |  |  |
| genetic reagent<br>(*D. melanogaster*) | *Ubi-Mer-Halo* | This study |  |  |
| genetic reagent<br>(*D. melanogaster*) | *Hpo-YFP* | (Su et al., 2017) |  |  |
| genetic reagent<br>(*D. melanogaster*) | *Halo-Snap-aPKC* | (Erdmann et al., 2019) |  |  |
| genetic reagent<br>(*D. melanogaster*) | *UAS-Crb RNAi* | Bloomington Drosophila Stock Center | BL35520 |  |
| genetic reagent<br>(*D. melanogaster*) | *UAS-Patj RNAi* | Bloomington Drosophila Stock Center | BL35747 | Validated in 10.1083/jcb.201611196 |
| genetic reagent<br>(*D. melanogaster*) | *UAS-Sdt RNAi* | Bloomington Drosophila Stock Center | BL33909 | Validated in 10.1083/jcb.201611196 |
| genetic reagent<br>(*D. melanogaster*) | *UAS-Ex RNAi* | Vienna<br>Drosophila<br>Resource Center | VDRC<br>109281 | Validated in<br>10.1016/j.de<br>vcel.2017.02<br>.004 |
| genetic reagent<br>(*D. melanogaster*) | *UAS-Utr-GFP* | Bloomington<br>Drosophila Stock<br>Center | BL34874 |  |
| genetic reagent<br>(*D. melanogaster*) | *aPKC<sup>as4</sup>* | (Hannaford et al.,<br>2019) |  |  |
| antibody | anti-PKC,<br>rabbit<br>polyclonal | Santa Cruz<br>Biotechnology | Lot#:<br>G2304 | tissue staining<br>(1:1000) |
| antibody | anti-Crb,<br>mouse<br>monoclonal | DSHB | Cq4 | tissue staining<br>(1:1000) |
| antibody | anti-FLAG,<br>mouse<br>monoclonal | Sigma Aldrich | Cat#F1804;<br>RRID:AB_262044 | S2 cell<br>staining (1:<br>20,000) |
| antibody | anti-Myc,<br>mouse<br>monoclonal | Cell Signaling | Product<br>#2276 | S2 cell<br>staining<br>(1:10,000) |
| Chemical | 1-NA-PP1 | Cayman | Cat.# CAYM-10954-1 |  |
| Chemical | Jasplakinolide | Cayman |  |  |

### Cortical Kib recruitment with Crb, Par6, aPKC in S2 cells

For experiments in Fig. 5, the following constructs were used: pMT-Kib-GFP, pMT-Par-6-Myc, pMT-aPKC, and pAc.5.1-Crb^i^-3xFLAG. Briefly, 2.7 x 10^6^ S2 cells (S2-DGRC) were transfected with total of 500ng of the indicated DNA constructs using dimethyldioctadecylammonium bromide (Sigma; Han, 1996) at 250µg/ml in 6-well plates. To induce expression of pMT constructs, 350µM CuSO4 was added to the wells 48h after transfection. Cells were collected 72h after transfection, fixed with 2% PFA/PBS for 15min, stained in PBS/0.1% Saponin/1% Normal goat serum (PSN) for 1h with primary (anti-Myc and anti-aPKC, see Table 1) and 1h with secondary antibodies, with 30 min washes. Cells were mounted in Prolong Diamond Antifade Mountant (Thermo Fisher Scientific) and cortical Kib localization was scored blind (at least 50 cells per condition) via widefield fluorescence using a Zeiss Axioplan 2ie microscope. Representative images in Fig. 5 were acquired using Zeiss LSM880 confocal microscope.

### Cortical Kib recruitment with Ed:GFP:aPKC in S2 cells

For experiments in Fig. 6, the following constructs were used: pMT-Ed:GFP or pMT-Ed:GFP:aPKC, Ubi>Kib-GFP-FLAG, Ubi>Kib^ΔaPKC^-GFP-FLAG, and Ubi>Kib^858-1288^-GFP-FLAG. Cells were transfected and expression of pMT constructs was induced as described above. Cells were resuspended 72h after transfection and gently mixed (in a 6-well plate) on an orbital shaker at 70 rpm for 30min to promote cell-cell clustering. After mixing, cells were collected, stained with primary (anti-aPKC and anti-FLAG, see Table 1) and secondary antibodies, mounted, and scored blind as described above.

