## Supplemental figures and legends for "Apical polarity and actomyosin dynamics regulate Hippo signaling by controlling Kibra subcellular localization"

### Figure S1

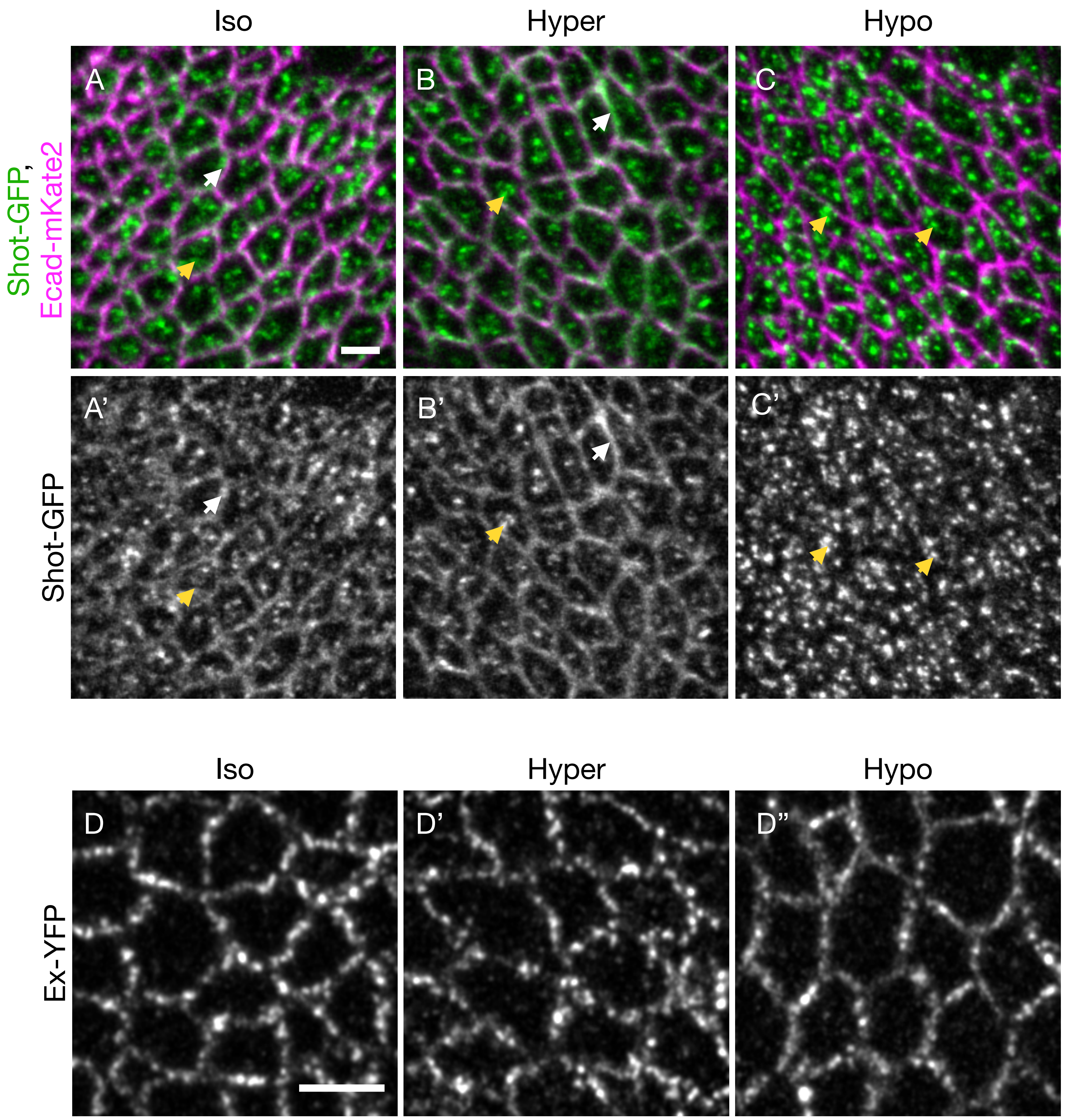

Iso
Hyper
Hypo
A
B
C
Shot-GFP, Ecad-mKate2
A'
B'
C'
Shot-GFP
Iso
Hyper
Hypo
D
D'
D''
Ex-YFP

### Figure S2

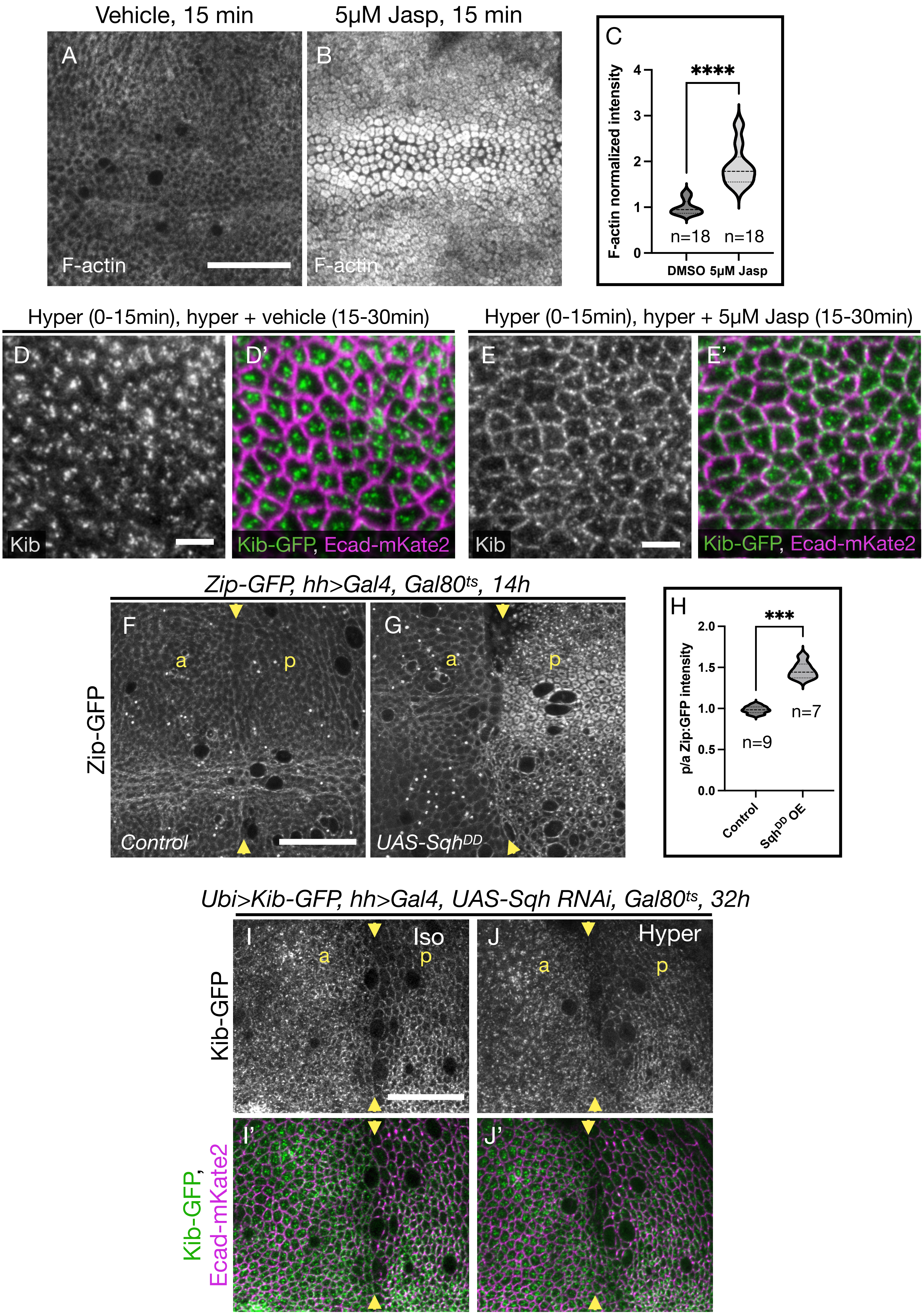

Vehicle, 15 min
5μM Jasp, 15 min
A
B
C
F-actin
F-actin
F-actin normalized intensity
****
n=18
n=18
DMSO
5μM Jasp
Hyper (0-15min), hyper + vehicle (15-30min)
Hyper (0-15min), hyper + 5μM Jasp (15-30min)
D
D'
E
E'
Kib
Kib-GFP, Ecad-mKate2
Kib
Kib-GFP, Ecad-mKate2
Zip-GFP, hh>Gal4, Gal80ts, 14h
Zip-GFP
F
G
a
p
a
p
Control
UAS-SqhDD
H
***
p/a Zip:GFP intensity
n=9
n=7
Control
SqhDD OE
Ubi>Kib-GFP, hh>Gal4, UAS-Sqh RNAi, Gal80ts, 32h
Kib-GFP
Kib-GFP, Ecad-mKate2
I
Iso
J
Hyper
a
p
a
p
I'
J'

### Figure S3

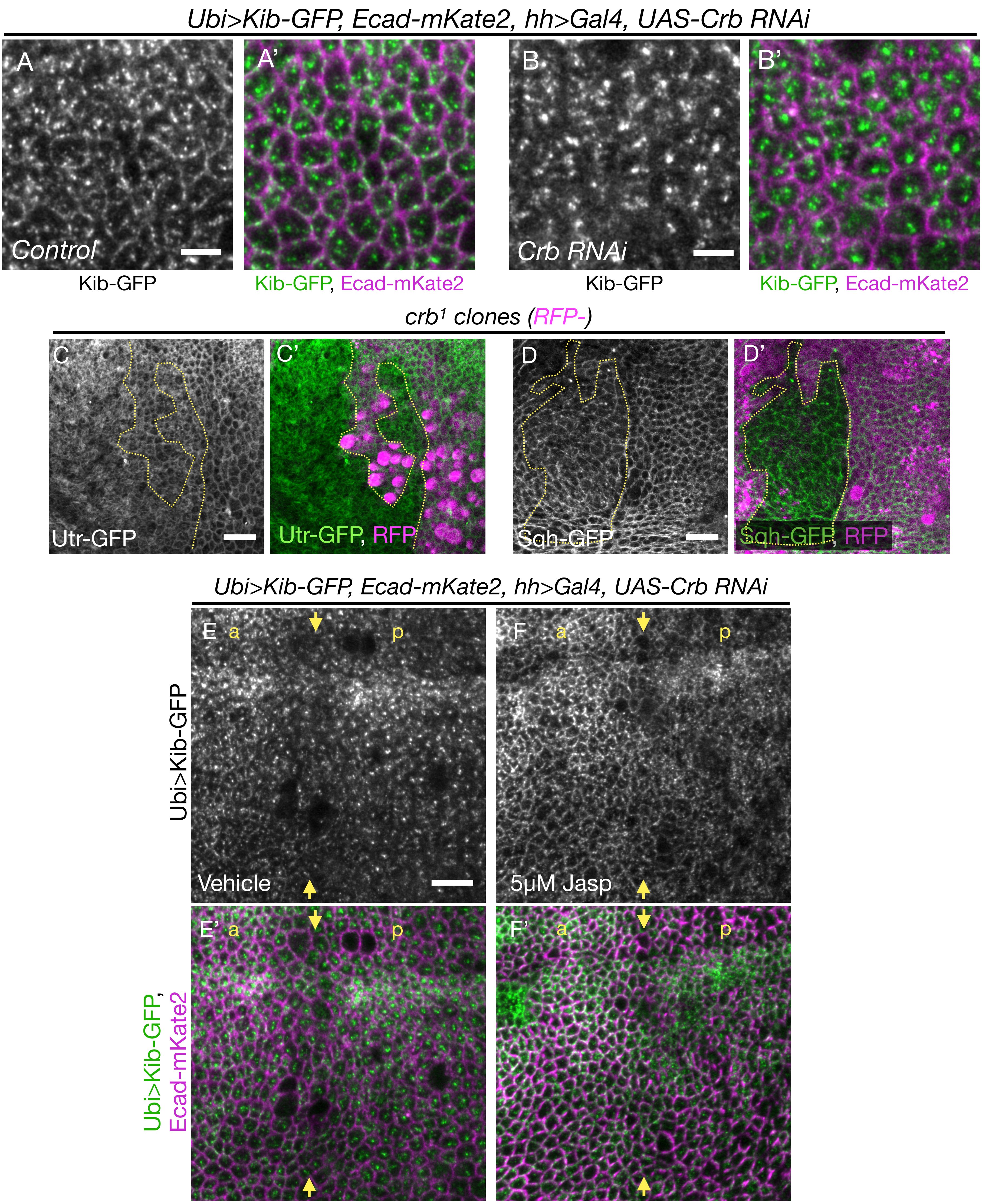

Ubi>Kib-GFP, Ecad-mKate2, hh>Gal4, UAS-Crb RNAi
A
A'
B
B'
Control
Crb RNAi
Kib-GFP
Kib-GFP, Ecad-mKate2
Kib-GFP
Kib-GFP, Ecad-mKate2
crb1 clones (RFP-)
C
C'
D
D'
Utr-GFP
Utr-GFP, RFP
Sqh-GFP
Sqh-GFP, RFP
Ubi>Kib-GFP, Ecad-mKate2, hh>Gal4, UAS-Crb RNAi
E
a
p
F
a
p
Ubi>Kib-GFP
Vehicle
5μM Jasp
E'
a
p
F'
a
p
Ubi>Kib-GFP,
Ecad-mKate2

### Figure S4

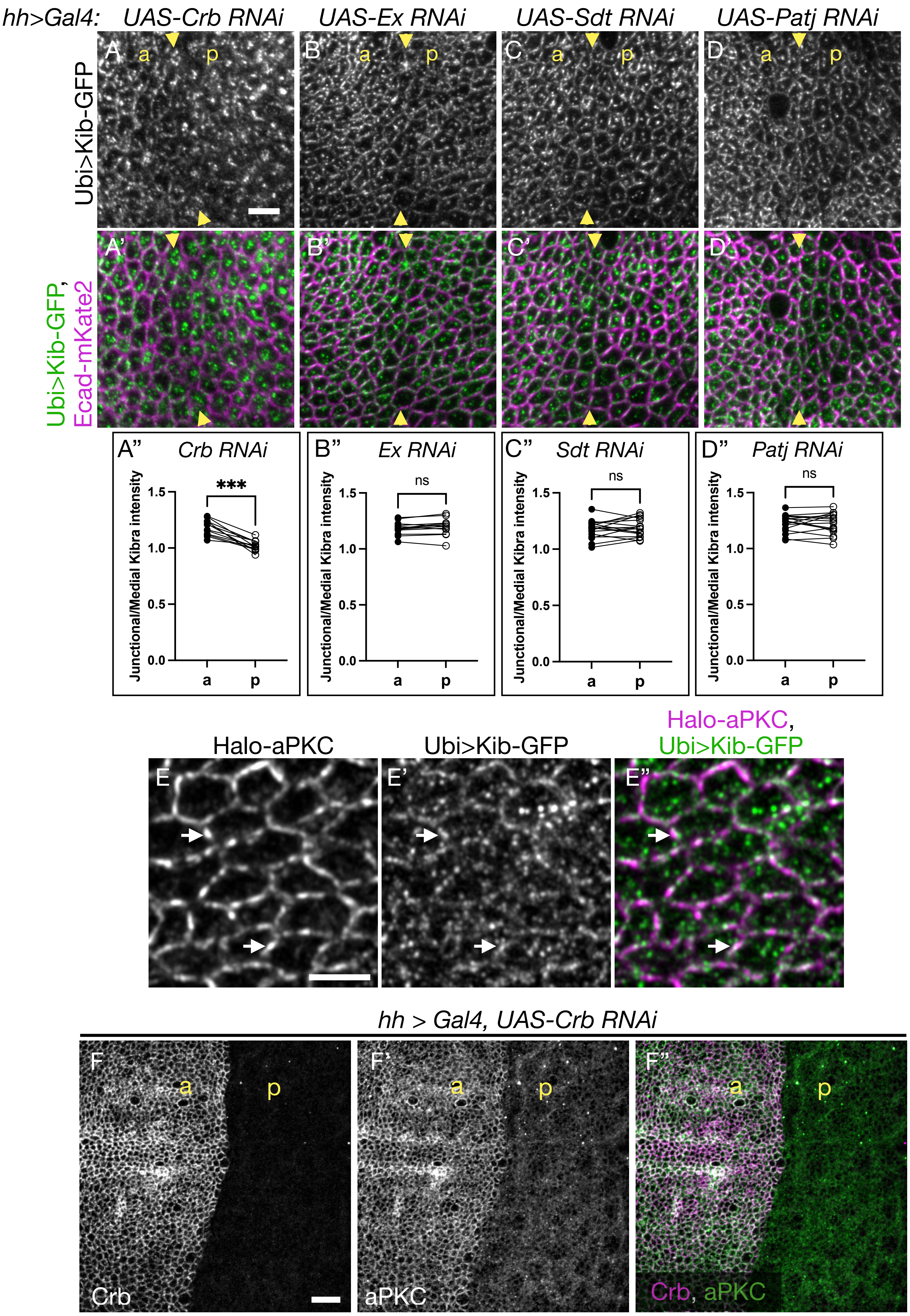

hh>Gal4: UAS-Crb RNAi
UAS-Ex RNAi
UAS-Sdt RNAi
UAS-Patj RNAi
Ubi>Kib-GFP
Ubi>Kib-GFP, Ecad-mKate2
A a p
B a p
C a p
D a p
A'
B'
C'
D'
A" Crb RNAi
B" Ex RNAi
C" Sdt RNAi
D" Patj RNAi
Junctional/Medial Kibra intensity
***
ns
1.5
1.0
0.5
0.0
a p
Halo-aPKC
Ubi>Kib-GFP
Halo-aPKC, Ubi>Kib-GFP
E
E'
E"
hh > Gal4, UAS-Crb RNAi
F a p
F' a p
F" a p
Crb
aPKC
Crb, aPKC

### Figure S5

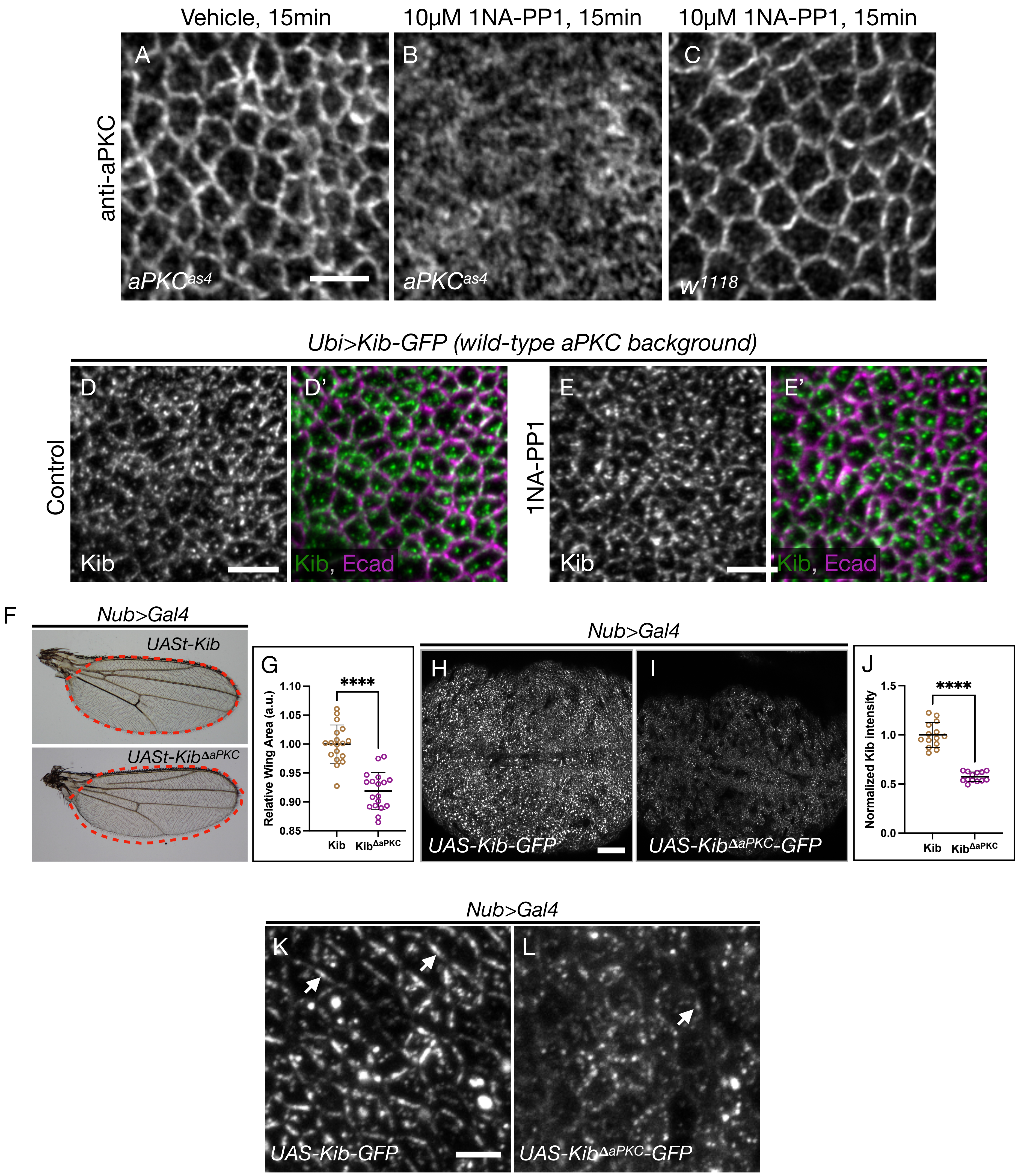

Vehicle, 15min
10μM 1NA-PP1, 15min
10μM 1NA-PP1, 15min
anti-aPKC
A
B
C
aPKCas4
aPKCas4
w1118
Ubi>Kib-GFP (wild-type aPKC background)
Control
D
D'
Kib
Kib, Ecad
1NA-PP1
E
E'
Kib
Kib, Ecad
F
Nub>Gal4
UASt-Kib
UASt-KibΔaPKC
G
Relative Wing Area (a.u.)
1.10
1.05
1.00
0.95
0.90
0.85
****
Kib
KibΔaPKC
Nub>Gal4
H
I
UAS-Kib-GFP
UAS-KibΔaPKC-GFP
J
Normalized Kib intensity
1.5
1.0
0.5
0.0
****
Kib
KibΔaPKC
Nub>Gal4
K
L
UAS-Kib-GFP
UAS-KibΔaPKC-GFP

### Figure S6

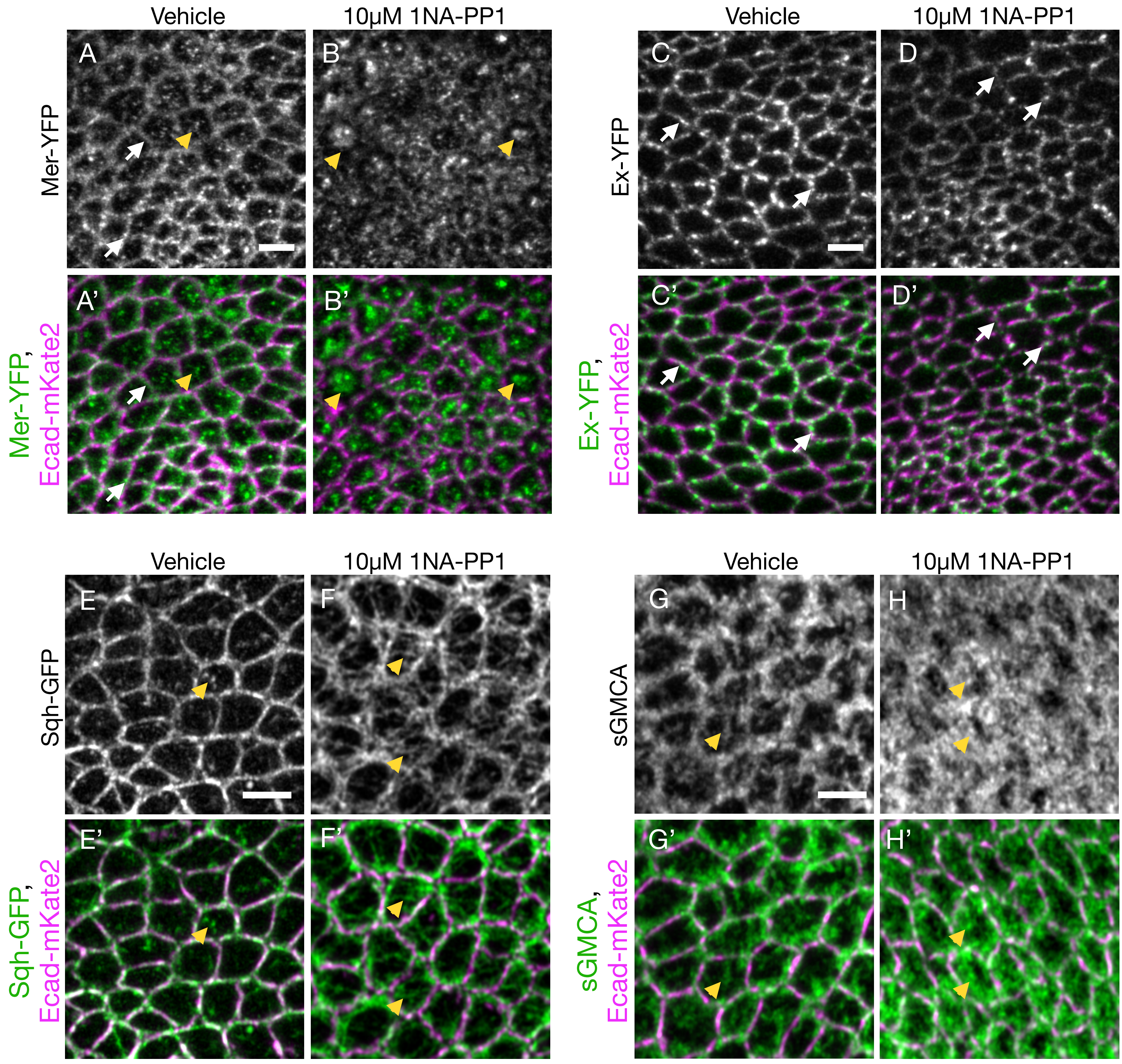

Vehicle
10µM 1NA-PP1
Vehicle
10µM 1NA-PP1
A
B
C
D
Mer-YFP
Ex-YFP
A'
B'
C'
D'
Mer-YFP,
Ecad-mKate2
Ex-YFP,
Ecad-mKate2
Vehicle
10µM 1NA-PP1
Vehicle
10µM 1NA-PP1
E
F
G
H
Sqh-GFP
sGMCA
E'
F'
G'
H'
Sqh-GFP,
Ecad-mKate2
sGMCA,
Ecad-mKate2

**Figure S1: The effect of osmotic shifts on Kib localization is specific.**
A-C’) Similar to Kib, Shot localizes at the junctional (white arrow) and medial cortex (yellow arrowhead) under isotonic conditions (A-A’). However, Shot remains both junctional and medial under hypertonic conditions (B-B’) and becomes predominantly medial under hypotonic conditions (C-C’).
D-D’’) Unlike Kib, Ex remains mostly junctional under osmotic shifts. Scale bars = 3µm.

**Figure S2: Medial Kib accumulation is mediated via actomyosin dynamics.**
A-B) Compared to control tissues (A), F-actin (sGMCA) was more stabilized in tissues treated with 5µM Jasp (B). Scale bar = 20µm.
C) Quantification of mean sGMCA intensity in tissues treated with DMSO or 5µM Jasp. Statistical significance was calculated using Mann-Whitney test.
D-E') F-actin dynamics are required to maintain Kib at the medial cortex. Tissues were first incubated under plain hypertonic conditions for 15 min to concentrate Kib medially, and then transferred to a hypertonic medium with DMSO (D-D') or 5µM Jasp (E-E'). Treatment with 5µM Jasp reverses medial Kib accumulation (E-E'). Scale bar = 5µm.
F-G) Compared to control tissues (F), transient ectopic expression of $Sqh^{DD}$ in the posterior compartment of the wing disc results in significant stabilization of the myosin heavy chain, Zip-GFP (G). Yellow arrows indicate the anterior-posterior (a-p) boundary. Scale bar = 20µm.
H) Quantification of the posterior/anterior (p/a) mean Zip:GFP intensity in control tissues and tissues transiently expressing $Sqh^{DD}$ in the posterior compartment. Statistical significance was calculated using Mann-Whitney test.
I-J') Transient depletion of Sqh in the posterior compartment of the wing imaginal disc leads to more junctional Kib (I-I') and prevents medial Kib accumulation under hypertonic conditions (J-J'). Scale bar = 20µm.

**Figure S3. Crb regulates Kib localization via actomyosin dynamics and junctional tethering.**

A-B') Kib is junctional and medial in control cells (A-A') but is more medial in cells depleted of Crb (B-B'). Scale bar = 5µm.

C-C') Apical F-actin organization is more medial in *crb*$^{1}$ somatic mosaic clones.

D-D') Junctional MyoII organization is disrupted in *crb*$^{1}$ somatic mosaic clones. Dashed yellow line in A-B' marks the clone (RFP negative) boundary. Scale Bar = 10µm.

E-F') While Kib accumulates medially upon depletion of Crb in the posterior (p) compartment of the wing imaginal disc (E-E'), this accumulation is blocked by Jasp treatment (F-F'). Note that while Jasp treatment enhances junctional Kib localization in the anterior (a) compartment, it fails to restore junctional Kib in the posterior, where Crb is depleted (F-F'). Scale Bar = 10µm.

**Figure S4. Crb does not control Kib localization via Ex, Sdt, or Patj.**
A-D'') While loss of Crb results in more medial Kib (A-A''), loss of Ex (B-B''), Sdt (C-C''), or Patj (D-D'') has no significant effect on Kib localization. Yellow arrowheads indicate the anterior-posterior (a-p) boundary. Scale bar = 5μm.
Statistical significance was calculated using Wilcoxon matched-pairs signed rank test.
E-E'') An apical projection of the wing imaginal disc cells showing that aPKC is highly enriched at the junctional cortex and that some junctional aPKC foci co-localize with Kib (white arrows). Scale bar = 3μm.
F-F'') Crb depletion in the posterior (p) compartment of the wing imaginal disc leads to significant decrease in cortical aPKC. Scale bar = 10μm.

**Figure S5. aPKC-mediated regulation of Kib in vivo.**
A-C) Validation of the specificity of $aPKC^{as4}$ inhibition by 1NA-PP1 in the wing imaginal disc. In tissues homozygous for *aPKCas4* allele, aPKC displays normal cortical localization (A). Treatment with 1NA-PP1 severely disrupts aPKC cortical localization (B). In contrast, wild-type aPKC is not affected by 1NA-PP1 treatment (C). Scale bar = 5µm.
D-E') Treatment with 1NA-PP1 does not affect Kib localization in the wild-type *aPKC* background. Scale bars = 5µm.
F-G) Ectopic expression of $Kib^{\Delta aPKC}$ (*UASt>*$Kib^{\Delta aPKC}$*-GFP*) using a wing-specific driver (*Nub>Gal4)* leads to slightly smaller wings compared to the expression of wild-type Kib (*UASt>Kib-GFP*).
H-J) Wild-type Kib (H) ectopically expressed specifically in the wing pouch is more stable than $Kib^{\Delta aPKC}$ (I & J). Scale bar = 20µm. Transgenes in F-I are identically expressed.
Statistical significance in G & J was calculated using Mann-Whitney test.
K-L) Similar to wild type Kib-GFP, ectopically expressed $Kib^{\Delta aPKC}$-GFP still localizes at the junctional cortex, albeit less prominently. Note that brightness in L was artificially enhanced. Scale bar = 3µm.

**Figure S6. The effect of acute aPKC inhibition on Hippo pathway components and actomyosin organization.**
A-B') Compared to control conditions (A & A') where Mer displays junctional and medial localization, Mer becomes more medial under aPKC inhibition with 1NA-PP1 (B & B').
C-D') Unlike Kib and Mer, Ex remains junctional under aPKC inhibition with 1NA-PP1.
E-H') aPKC inhibition with 1NA-PP1 results in more medial myosin (E-F') and F-actin (G-H') organization. Wing imaginal tissues in A-H' are homozygous for *aPKCas4* allele. White arrows point to junctional cortex. Yellow arrowheads point to medial cortex. Scale bars = 3μm.
